# Circumventing glioblastoma resistance to temozolomide through optimal drug combinations designed by quantitative systems pharmacology and machine learning

**DOI:** 10.1101/2024.05.31.596811

**Authors:** Sergio Corridore, Maïté Verreault, Hugo Martin, Thibault Delobel, Cécile Carrère, Ahmed Idbaih, Annabelle Ballesta

## Abstract

Glioblastoma is currently associated to a dismal prognosis despite intensive treatment involving maximal-safe surgery, radiotherapy and temozolomide (TMZ)-based chemotherapy. Disease progression or relapse is often due to initial or acquired resistance to temozolomide, which may be mediated by the over-expression of the repair enzyme MGMT. To design TMZ-based drug combinations circumventing the initial resistance of MGMT-overexpressing cells, a quantitative systems pharmacology (QSP) model representing TMZ cellular pharmacokinetics-pharmacodynamics and their connection to the most altered pathways in GBM was developed. This digital network representation of TMZ cellular pharmacology successfully integrates, in a mechanistic fashion, multi-type time- and dose-resolved datasets, available in control or MGMT-overexpressing cells. *In silico* target inhibition screening identified an optimal antitumor strategy consisting in priming cancer cells with inhibitors of the base excision repair and of the homologous recombination pathway prior to TMZ exposure. This drug combination was validated in dedicated experiments, thus allowing to re-sensitize cells which were initially resistant to TMZ. Using machine learning, functional signatures of response to such optimal multiagent therapy were derived to assist decision making about administering it to other cancer cell lines or patients. The developed framework can be extended to account for additional patientspecific altered pathways and may be translated towards the clinics by representing the tumor micro-environment and drug whole-body pharmacokinetics. Overall, we successfully demonstrated the relevance of combined QSP and machine learning to design multi-agent pharmacotherapies circumventing initial tumor resistance.

**One Sentence Summary:** An integrated *in vitro*-*in silico* approach allowed to design optimal drug combinations re-sensitizing temozolomide-resistant glioblastoma cells.

## INTRODUCTION

As science helps us discover that cancer diseases are multiple and patient-specific, great efforts are made towards cancer treatment personalization. Personalized cancer management offers to dramatically improve patient responses by integrating multi-type patient-specific datasets into treatment decision. Recent technological breakthroughs currently allow for the collection of multiple preclinical and clinical high-throughput data, and decision-making algorithms offering to translate such multi-level knowledge into patient-specific therapies are critically needed. The complexity inherent to the integration of multi-type and multi-scale datasets appeals for mathematical and computational approaches providing *in silico* representation of the patient and its disease in the shape of a *digital twin* (*1–3*).

The field of Quantitative Systems Pharmacology (QSP) is dedicated to the development of such theoretical approaches with the specificity of designing a *mechanism-based* representation of the patient’s tumor accounting for key alterations of molecular pathways involved in drug pharmacokinetics (PK), pharmacodynamics (PD), and antitumor efficacy (*4–6*). Indeed, anticancer drug toxicity and efficacy are ultimately determined at the molecular scale by the response of gene and protein networks involved in the drug response in different cell populations. Hence, theoretical models of cell type-specific regulatory pathways constitute a reliable physiological basis from which the treatment can be optimized. The tumor model may further be included into a patient model to account for whole-body drug PK or dose-limiting toxicities. Thus, such theoretical models do not subdivide living organisms into independent components, but rather, recognize that genes, proteins, cells, and organs interact with each other and with the environment in complex ways that can vary over time. QSP studies may inform on the strength of a hypothesis, extrapolating preclinical and clinical findings. One successful example of QSP studies to inform precision drug dosing is shown for nivolumab in the context of melanoma, non-small cell lung cancer, and renal cell carcinoma. A QSP model composed of a population PK model and of a PD-1/ligand receptor binding model, integrated both clinical and *in vitro* data and supported approval of nivolumab 480 mg every 4 weeks a dosing regimen with less frequent administration as compared to the initial one (*7*). Recently, QSP models have been used to generate virtual patient populations (*8*) which can further be studied through machine learning to inform on key model parameters driving drug efficacy and to design signature of response (*5*). Here, we present a QSP modelling framework that was developed for temozolomide (TMZ), an alkylating agent largely utilized against glioblastoma (GBM), the most frequent and aggressive primary brain cancer in adults.

Despite very intensive treatments combining maximal safe surgical resection, radiation therapy (RT) and TMZ-based chemotherapy, the prognosis of GBM patients remains poor with a median overall survival below 18 months (*9*). Even though GBM represents 70% of all adult primary malignant brain tumors, its incidence remains relatively low, around 3 per 100,000 individuals (*9*). As a result, patient recruitment for classical clinical trials may be challenging which impairs classical drug development and innovative approaches prioritizing hypothesis/therapies to test in the clinics and enabling more personalized strategies according to QSP-based biomarkers would be particularly helpful (*10*). The cornerstone of GBM pharmacotherapies, TMZ, usually demonstrates moderate efficacy when administered as a single agent, in combination with RT, which may be explained by the intrinsic resistance to chemotherapy of some GBM cell subpopulations or by their capacity to evolve under treatment pressure towards resistant cell phenotypes (*11, 12*). The heterogeneous molecular profiles of GBM, together with the success of combining cytotoxics to targeted therapies in other cancers, suggest that TMZ-based combination therapies addressing more than one oncogenic pathway would offer better efficacy for patients (*13*). Thus, we focused here on optimizing the combination of TMZ with targeted agents affecting key molecular pathways.

Several high-throughput and/or mechanistic studies have shed light on GBM genetic alterations and main mechanisms of TMZ resistance which we accounted for in our study. According to an integrated analyses of multi-dimensional genomic data from The Cancer Genome Atlas (TCGA), GBM most frequent genetic alterations occurs in the RB of the cell cycle, p53, and RTK/RAS/ PI(3)K receptor tyrosine kinase pathways (*14*). TMZ is an alkylating agent that undergoes a two-step activation process leading to the formation of several types of DNA adducts. Among them, O6-meG is the less frequent but the most cytotoxic lesion if not repaired by O6-methylguanine-DNA methyl-transferase (MGMT) (*15, 16*). The presence of O6-meG adducts triggers futile loops involving mismatch repair (MMR), that produce DNA double stranded breaks and activate the ATR-CHK1 axis and P53 network towards cell cycle arrest and possibly cell apoptosis (*17*). The methylation status of *MGMT* gene promoter is an important molecular biomarker of TMZ response (*18*). A recent CRISPR-Cas9 whole-genome screen in GBM patient-derived stem cells identified the core members of the MMR pathway MLH1, MSH2, MSH6, and PMS2 as the only four genes associated to resistance to high-dose TMZ while multiple DNA repair pathways including HR or Fanconi anemia/interstrand crosslink repair were identified as intrinsic resistance mechanisms by exposing cells to low-dose TMZ (*19*).

Here, we present a QSP representation of TMZ PK-PD and key regulatory protein networks involved in the drug response. It was developed through an iterative approach, involving the integration of numerous multi-type datasets (Fig. 1). The model was calibrated to represent a sensitive and a MGMT-overexpressing resistant GBM cell line, mimicking a heterogeneous tumor with two clones. Once validated, those cell-type specific models were utilized to design optimal TMZ-based drug combinations to circumvent TMZ resistance. The model-derived optimal drug combination was validated in dedicated experiments which successfully demonstrated its efficacy and superiority as compared to other tested pharmacotherapies in both cell types. Next, we used machine learning classification to derive a signature for responders of this optimal drug combinations.

**Figure 1.**
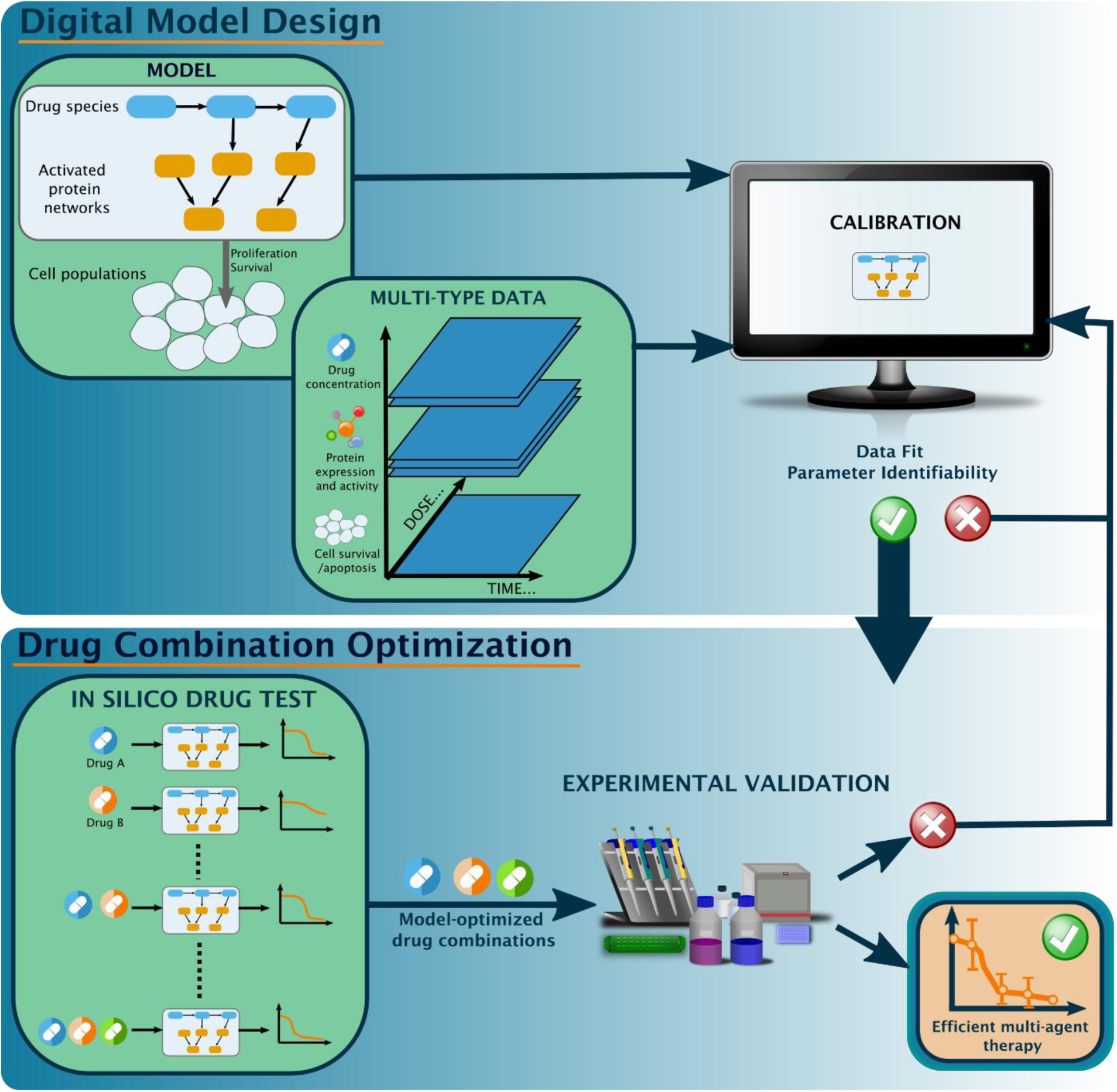
Workflow of systems pharmacology approach for the design of optimal drug combinations.

## RESULTS

### TMZ cellular pharmacokinetics-pharmacodynamics model

A physiologically-based model of TMZ cellular PK-PD was designed through an iterative process of data integration (Fig. 1). It is composed of a previously published model of TMZ PK (*20*) and a mechanistic PD model representing TMZ-induced DNA damage, DNA damage repair and ATR-mediated sensing, cell cycle, p53 pathways and apoptosis, such pathways being among the most disrupted in TCGA GBM patients (Fig. 2, Supplementary Materials (*14*)). Regarding the drug PK, the model represents TMZ activation into its metabolite MTIC which subsequently decomposes into AIC—an inactive metabolite—and a methyldiazonium cation, the DNA-methylating species, those two reactions being pH-dependent. TMZ and AIC cellular uptake and efflux are also included, MTIC being assumed not to cross the cell membrane.

**Figure 2.**
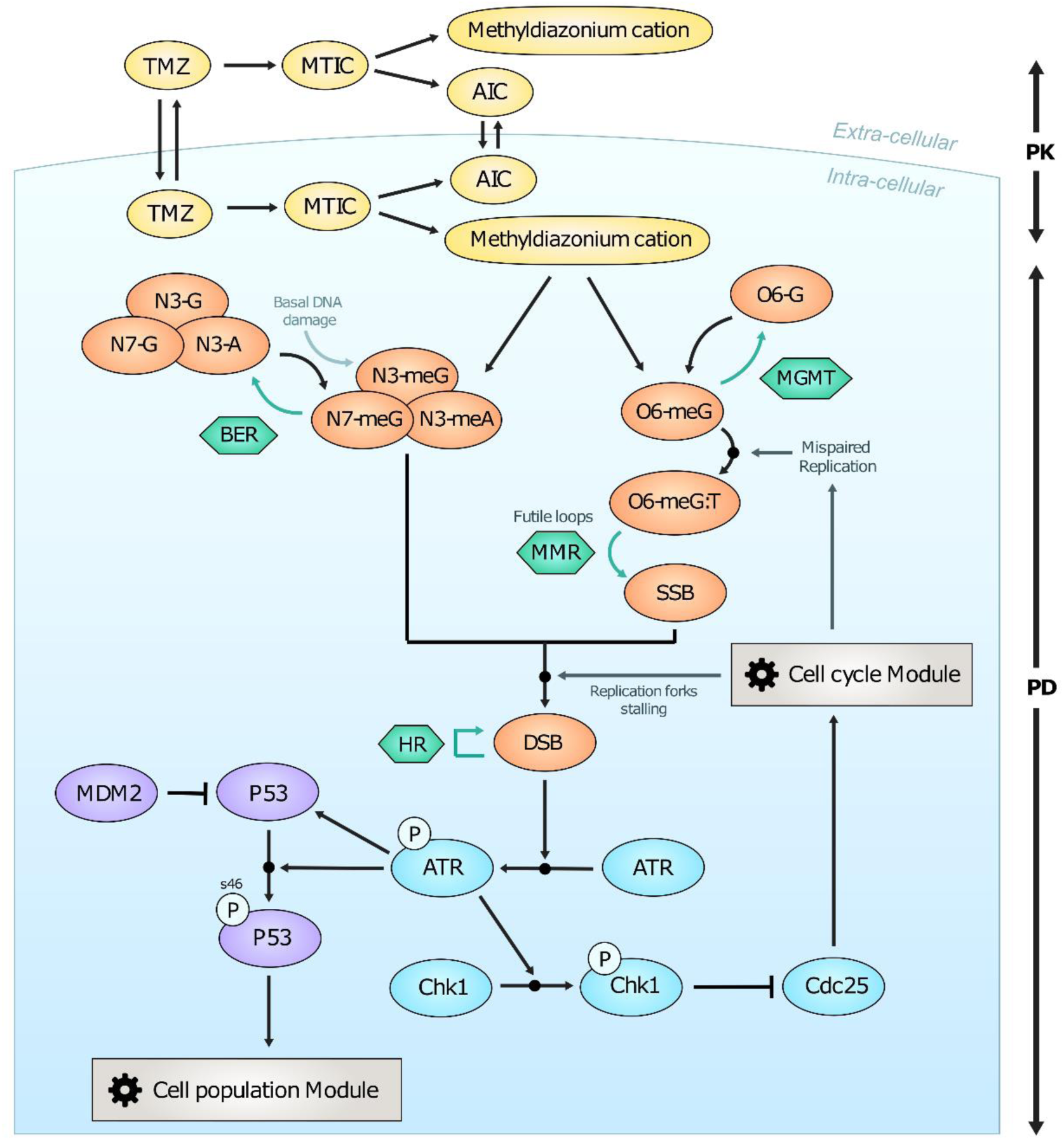
TMZ cellular PK-PD model. Color code is as follows: yellow for drug PK, orange for DNA damage, green for DNA repair mechanisms, blue for ATR-CHK1 axis, and purple for P53 network.

TMZ PD is driven by the methyldiazonium cation that forms four types of adducts on the DNA: N7-methylguanine (N7-meG), N3-methylguanine (N3-meG), N3-methyladenine (N3-meA) and O6-methylguanine (O6-meG) adducts. N7-meG, N3-meG and N3-meA adducts (lumped into the N37-meAG variable) may be repaired by BER whereas O6-meGs are specifically repaired by MGMT. The O6-meG:C pairs of bases are mispaired during replication and O6-meG:T are created thus triggering the MMR system which excises the T base but immediately re-inserts another one due to the mispair properties of O6-meGs. Those futile loops result in persistent Single Stranded Breaks (SSB) on the DNA, which may induce DNA lesions through cell-cycle dependent mechanisms as they may collide with replication forks leading to Double Stranded Breaks (*DSBs*). In turn, DSBs are repaired by homologous recombination (HR). As the only needed cell cycle input lays in the information of the cell being or not in S-phase, the cell cycle module is 24h-periodic binary variable representing a S-phase marker (Supplementary Information). We also considered a basal formation of DNA damage due to TMZ-independent mechanisms occurring in control conditions, which are counted in the N37-meAG adducts for the sake of simplicity, and which are thus repaired by BER.

The presence of DSBs triggers ATR kinases phosphorylation. Once activated, ATR phosphorylate CHK1 protein which enhances the degradation of CDC25, leading to cell cycle arrest (*21*). In parallel, ATR activates the P53 network including the P53-MDM2 negative feedback loop is activated and P53 phosphorylation at Ser46 leads to the launch of apoptosis (*22*). Such intracellular PK-PD model was then connected to a cell population module which describes the dynamics of a homogenous cell population during TMZ exposure. Cell death rate depends on the cumulative amount of phosphorylated P53 such that a cell is supposed to go to apoptosis when it passes a critical time-dependent threshold, as suggested by single cell experimental data (Supplementary Information (*23*)).

### Model calibration

The TMZ cellular PK-PD model counts 22 state variables and 66 parameters, presented in Table S1 and S2, that were meticulously estimated to obtain model behaviors accurately mimicking dose- and time-dependent experimental observations (Supplementary Information, Section 3). We performed a parameter estimation strategy consisting of two steps: *i*) *step 1,* in which a subset of 37 parameters were independently evaluated thanks to the availability of quantitative information in the literature (n=25) or fixed to realistic values in agreement with biological knowledge (n=12), *ii*) *step 2,* consisting in the estimation of the 32 remaining parameters based on extensive experimental studies performed in two populations of LN229 human GBM cell lines: the control (MGMT-) and MGMT-overexpressing (MGMT+) ones. This step integrated the data of four studies documenting DNA adducts, DSBs, total and phosphorylated CHK1, CDC25, total and phosphorylated P53, cell death and survival during TMZ exposure in a time- and/or dose-dependent manner (approx. 300 datapoints, Supplementary Information Section 3, (*24–27*)). Six parameters were inferred from the other ones through algebraic equations derived from the assumption that the system was at steady state at initial conditions (Supplementary Information, Section 2). The remaining 25 parameters were estimated through an optimization problem aiming to fit the above-mentioned datasets (see Methods).

The best-fit model accurately represented all datasets both with respect to TMZ dose (Fig. 3) and to temporal dynamics (Fig. 4), reflecting the different behaviors of sensitive LN229 MGMT- and resistant MGMT+ cell lines. Experimentally, both O6-meG and DSB amounts increased faster along with TMZ dose in the sensitive cell line as compared to the resistant one, such behavior being reproduced by the model (Fig. 3(**A**-**B**)). The model slightly underestimated both O6-meG and DSB amounts in MGMT+ cells, which suggested that possible saturation mechanisms may occur for high TMZ doses, which were not included in the model. Next, P53 and P53-Ser46 amounts rapidly reached saturation when TMZ dose was increased, and this level increased with exposure time in MGMT-cell experiments (Fig. 3(**C**-**D**)). Such phenomenon was caught by the model which further predicted low amounts of P53 species in MGMT+ cells. In MGMT-cell line, both the experiments and the model showed that the percentage of apoptotic cells was positively correlated with TMZ dose and exposure duration, reaching a maximum of approx. 40% after 5 days of exposure at concentrations greater than 75 μM (Fig. 3(**E**)). On the opposite, cell death in MGMT+ condition never exceeded 8%, for any dose smaller than 125 μM and any duration below 6 days. Finally, MGMT-cell survival data showed a pronounced sensitivity to TMZ with an IC50 of 5 μM. For MGMT+, cells were largely resistant to TMZ for concentration below 256 μM, and the IC50 equal to 400 μM (Fig. 3(**F**)). All those dynamics were quantitatively reproduced by the model, which gives a good estimation of IC50 for both cell lines, in particular 8 μM and 450 μM, respectively for MGMT- and MGMT+.

**Figure 3.**
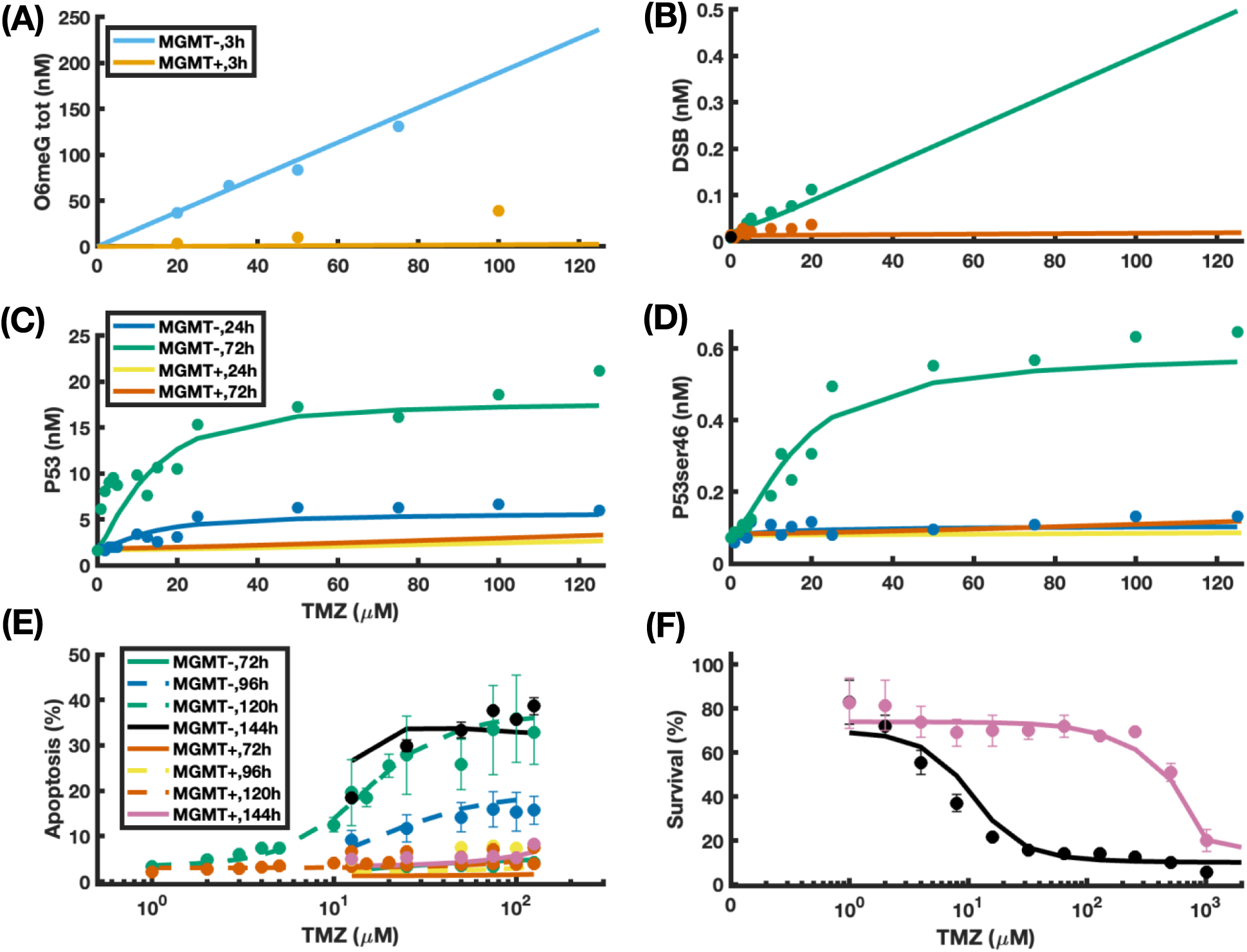
Dose-dependent TMZ PK-PD and efficacy in MGMT- and MGMT+ cell lines. Dots and bars represent mean and standard deviation of experimental results for different TMZ doses, when available. Lines are the best-fit model.

**Figure 4:**
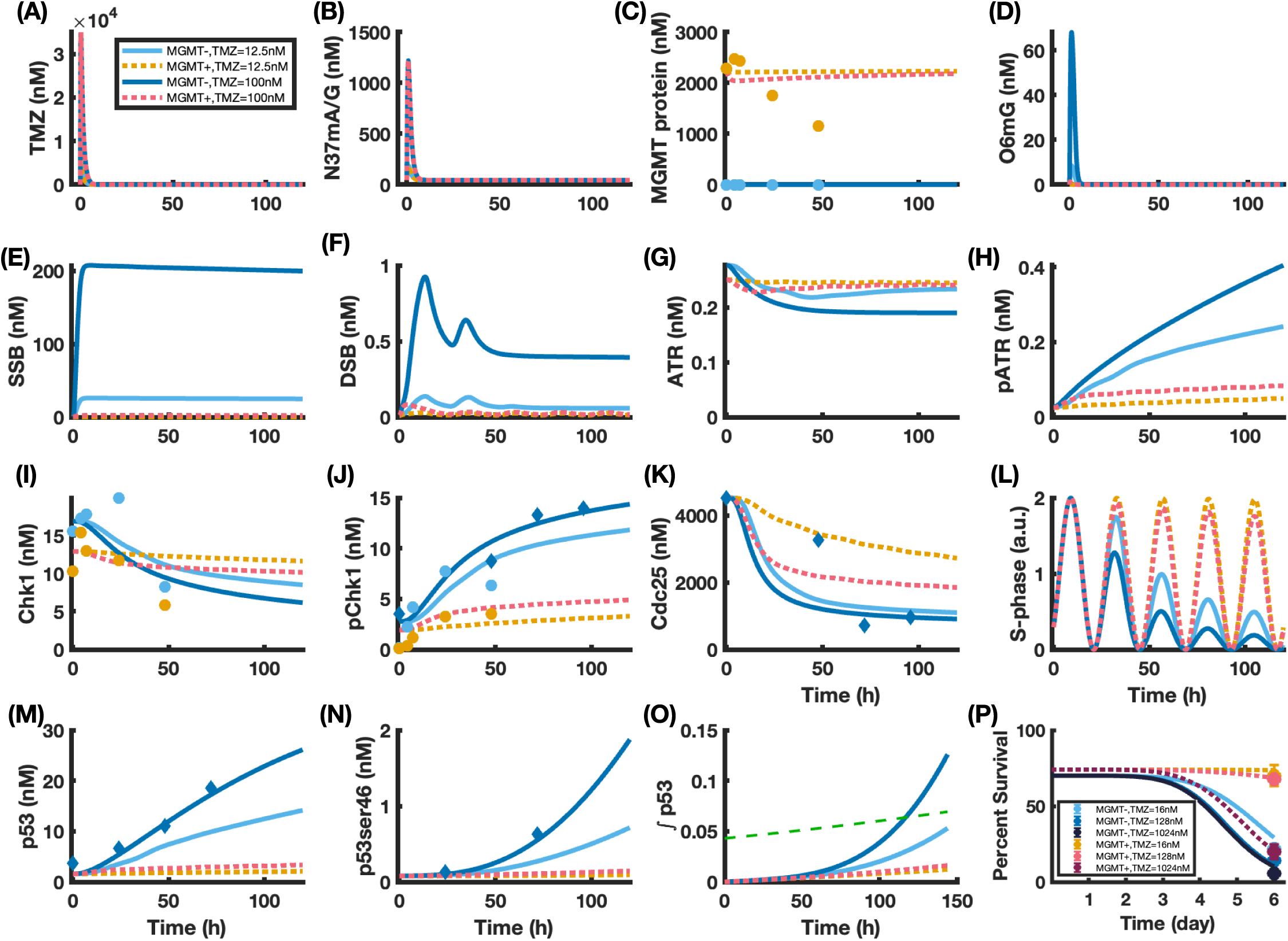
Time-dependent TMZ cellular PK-PD. TMZ low (12.5 μM) or high (100 μM) dose conditions are shown for MGMT-(respectively cyan and blue) and for MGMT+ (respectively orange and red) cells. Dots represent experimental results for different exposure durations, when available. Lines are the best-fit model. **(A)** Intracellular TMZ concentration, **(B, D-F)** TMZ-induced DNA damage concentration, **(C)** MGMT protein concentration, **(G-J)** Unphosphorylated or phosphorylated ATR and CHK1 concentrations, **(K)** Cdc25 concentration, **(L)** S-phase marker time profile, **(M-N)** Unphosphorylated and Ser46 phosphorylated P53 concentrations, **(O)** P53 cumulative amount (area under the curve), the green line indicates the threshold for apoptosis launching, **(P)** Cell survival over time for three TMZ doses (16, 128 and 1024 μM).

Next, the temporal evolution of the concentrations of drug species and proteins described in the model were investigated both *in vitro* and *in silico* for two TMZ doses, low (12.5 μM) and high (100 μM), for both MGMT- and MGMT+ cell lines (Fig. 4). In agreement with previously published PK modeling studies (*20*), TMZ intracellular concentration rapidly increases after exposure start, reaches its peak in less than 15 min, and then decreases due to its degradation into MTIC, in all conditions (Fig. 4(**A**), Fig. S1). N37-meAG concentration displayed the same fast dynamics as it achieves its peak approximately 1h after exposure start, in a dose-dependent manner, yet with the same dynamics for MGMT- and MGMT+ cells (Fig. 4(**B**)). O6meG dynamics is also rapid but in MGMT-cells, while this type of adducts is almost absent in MGMT+ cells for any TMZ concentration. This is respectively due to low or overexpressed MGMT protein amount in MGMT- and MGMT+ cells, as confirmed by the biological observations (Fig. 4(C-D)). Data of MGMT protein levels in MGMT+ cells after TMZ exposure at 12.5 μM are reproduced only in part by the best-fit model which underestimates MGMT decreases, suggesting a possible TMZ-independent degradation of the protein.

In MGMT-cells, the presence of O6-meG adducts produces G:T mispairs, both species displaying very rapid kinetics and disappearing in only few hours after TMZ exposure start, which explains the absence of cell cycle-mediated oscillations in these variables. The MMR futile loops then create SSBs which amount reaches its peak approximately 6h after exposure start and remains at a nearly constant value afterwards, with a slightly decreasing trend due to SSBs conversion into DSBs (Fig. 4(**E**) and Fig. S2(**E**)). SSB amounts are cell line- and dose-dependent as they are maximum for the high dose of TMZ in MGMT-cells, then low in MGMT-cells exposed to low dose or MGMT+ cells exposed to high dose and minimum for MGMT+ cells exposed to low dose. DSB level logically rises in the presence of SSBs and displays 24h-oscillations because of the effect of the cell cycle (Fig. 4(**F**), Fig. S2(**G**)). Indeed, DSB are only created when the S-phase marker is close to its peak indicating that replication mechanisms are ongoing (Fig. 4(**L**), Fig. S3(**F**)).

The presence of DSBs activates the ATR-CHK1 axis, which consists in the phosphorylation of ATR only once DSBs reach a specific threshold and, subsequently, the phosphorylation of CHK1 (Fig. 4(**G**-**J**), Fig. S3(**A**-**D**)). The best-fit model provides a good fit of phosphorylated CHK1 data in MGMT- and MGMT+ cell lines while only modestly agrees with total CHK1 experimental levels in MGMT+ cells. As confirmed by the biological observations, the increase in pCHK1 produces the degradation of CDC25 which arrests the cell cycle when reaching a critical threshold (Fig. 4(**K**, **L**), Fig. S3(**E**, **F**)). The cell cycle is gradually arrested over 6 days in MGMT-cells for both doses and in MGMT+ for highest one. On the opposite, in MGMT+ cells exposed at 12.5 μM, the cell cycle is not affected by TMZ. ATR phosphorylation further triggers the activation of the P53 pathway through both an increase in total P53 and in the activated fraction of P53 leading to cell apoptosis (P53ser46), as can be seen in MGMT-cells for both doses (Fig. 4(**M**, **N**), Fig. S4(**A**, **B**)). In parallel, the increase in both P53 and P53ser46 leads to the transcription of MDM2 mRNA and, therefore, to the translation of MDM2 protein, starting 2 days after TMZ exposure beginning (Fig. S4(**C**, **D**)). Consequently, MDM2 inhibits P53 expression which time profile changes of curvature when MDM2 protein becomes non-zero (Fig. S4(**A**, **D**)). The cumulative amount of P53 is then compared to a time-dependent threshold curve that corresponds to the cell state in which apoptosis is triggered leading to a decrease in cell survival (Fig. 4(**O**, **P**)).

### Parameter identifiability and machine learning-based sensitivity analysis

The accuracy of the best-fit model was further explored by assessing parameter identifiability and correlation. Parameter distributions displayed sharp profiles on the search domain and parameter coefficients of variations (CVs) ranged between 3 and 36.3%, thus demonstrating parameter practical identifiability (Fig. 5 (**A**) and Table S2). Moreover, as shown by the vine-copula multivariate analysis (see Methods), parameters were largely uncorrelated (Fig. 5 (**B**)). Three parameters related to ATR and P53 activation rates displayed correlation (k_ATR,_ K_ATR_ and kp_ser46_, pairwise R=0.72, −0.83, −0.71). This is likely to derive from the unavailability of ATR amount data and means that the current model may only inform on both sequential activation steps without being able to estimate the kinetics of each of them. In MGMT+ cells, the model could accurately account for the activity of the pool of MGMT proteins present in the cell but could not provide an estimation neither of the activity of a single MGMT protein nor of the absolute protein concentration, due to a lack of quantitative data informing these quantities separately (MGMT_0_ and k_MGMT_, R=-0.62), Next, we performed a parameter importance analysis to determine which molecular events drove the response to TMZ, using the area under the curve (AUC) of cell survival for different TMZ concentration as the output of interest (Fig. 5(**C**), see Methods). The most important parameters for both MGMT- and MGMT+ cell lines were the ones related to the cell apoptosis module (mainly UpAsy, tED50, Stiff), P53 translation and phosphorylation parameters (kf_P53_, k_P53_, K_ATR_, kp_ser46_), ATR activation rate (k_ATR_) and pATR degradation rate (kd_pATR_). In the MGMT+ cell lines only, high importance values were also assigned to TMZ PK parameters (p_T_, p_T2_) and to Homologous Recombination activity (k_HR_). Altogether, these results demonstrated that important parameters were distributed throughout the model thus confirming that all parts were necessary to accurately predict TMZ sensitivity.

**Figure 5.**
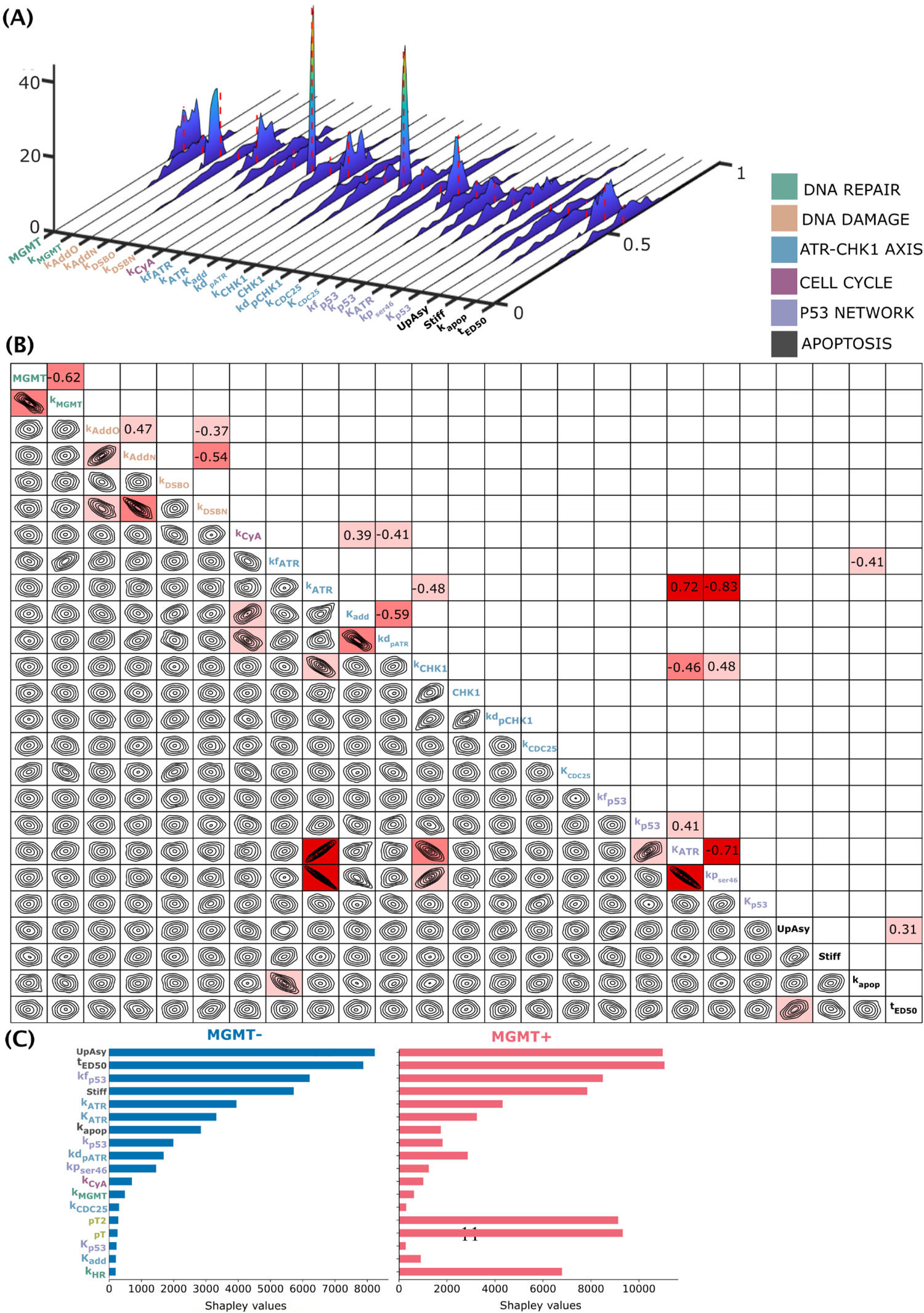
(next page): TMZ model parameter identifiability, correlation and importance. **(A)** Distribution of best-fit model parameters, **(B)** Correlation between model parameters as computed by fitting a vine copula model to best-fit parameter sets, **(C)** Parameter importance on TMZ dose-response AUC.

### Drug target identification and optimization of drug combinations

The calibrated TMZ model was then employed to identify optimal drug targets which inhibition would enhance TMZ antitumor efficacy in resistant cells. An *in silico* search was performed by inhibiting each possible target present in the model over the whole TMZ exposure duration. These scenarios thus assumed that the administration of the compound inhibiting the target occurred prior to TMZ exposure and that the inhibition last over the entire exposure duration, i.e. 5 days. The first step aimed at identifying drug combinations composed of TMZ and a single targeted molecule, thus only inhibiting one target at a time, at various levels from 0 to 100% corresponding to different target affinities or doses (Fig. 6(**A**)).

**Figure 6:**
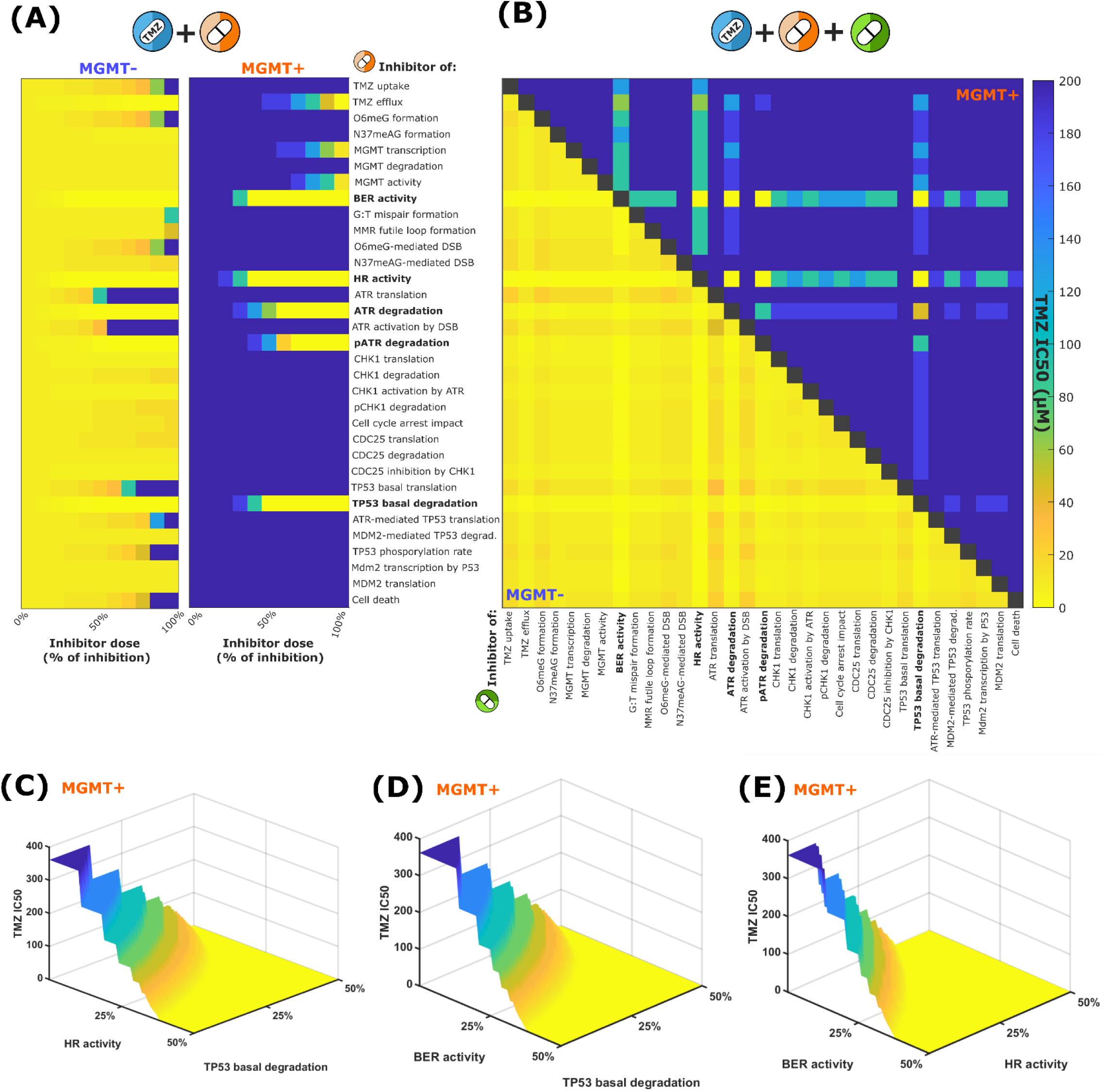
Drug target identification. **(A)** Investigation of efficacy of inhibition of a single target, at different levels, combined to TMZ exposure, in MGMT-sensitive (first column) and MGMT+ resistant (second column) cells, **(B)** Investigation of efficacy of inhibition of two targets at a level of 30%, combined to TMZ exposure, in MGMT-(lower triangle matrix) and MGMT+ resistant (upper triangle) cells, **(C)** TMZ IC50 with inhibitions of P53 degradation and HR activity at specified levels, **(D)** TMZ IC50 with inhibitions of P53 degradation and BER activity at specified levels, **(E)** TMZ IC50 with inhibitions of BER and HR activity at specified levels in MGMT+ cells.

Five target inhibitions prove efficient to increase TMZ activity in resistant cells: BER and HR repair activity; and ATR, pATR and P53 degradation. BER or HR inhibition at a level of 40% allowed to decrease TMZ IC50 to zero, thus meaning that the target inhibition by itself caused more than 50% cell death. P53, ATR and pATR degradation inhibition at respective levels of 50%, 60% and 70% achieved the same efficacy. Of note, the more intuitive solution consisting in inhibiting MGMT expression or activity was suboptimal as unrealistic inhibition levels close to 100% would have been required to significantly lower TMZ IC50 in resistant cells. Logically, these optimal drug combinations were also efficient in MGMT-cells (Fig. 6(**A**)).

Next, we explored pharmacotherapies combining TMZ with two targeted molecules to intend to further lower the levels of inhibition for each target separately. This may be equivalent to lower the inhibitor doses. We thus computed TMZ IC50 after moderate inhibition (i.e. 30%) of both targets (Fig. 6(**B**)). Only seven drug combinations led to a TMZ IC50 value of zero. They consisted to a combined inhibition of BER and HR repair activity or an inhibition of either of these pathways associated with an inhibition of ATR, pATR or P53 degradation. Regarding MGMT-cells, almost all drug triplets achieved high efficacy. We then further varied the inhibitors doses to search for the drug combinations associated with the lowest inhibition levels required for efficacy in resistant cells. Under this criterion, the combined inhibition of BER and HR pathways prior to TMZ administration was the best multi-agent strategy and was selected for further experimental validation (Fig. 6(**C-E**)).

### Experimental validation of optimal drug combinations

Once identified, the optimal TMZ-based drug combinations were tested in LN229 MGMT- and MGMT+ cell lines. Cells were exposed to all possible combinations of TMZ, niraparib, a PARP inhibitor that decreases BER repair efficiency, and RI-1, a HR inhibitor (Fig. 7(**A-C**)). As expected, the efficacy of TMZ given as a single agent was high in MGMT-sensitive cells and low in MGMT+ resistant cells (respective IC50=47 μM and 344 μM, t-test p= 0.0005), thus confirming the published data used for model calibration. Niraparib and RI-1 as single agents displayed similar cytotoxicity in MGMT- and MGMT+ cells with respective IC50 of 2.41 ±0.77 μM and of 1.83 ±0.15 μM for niraparib, and respective IC50 of 15.40 ±0.80 μM and of 15.22 ±0.23 μM for RI1, high variability being observed for RI-1 (Fig. S6). Based on those results, drug concentrations reducing cell viability by 20% for both cell lines were selected for combinations with TMZ and thus set to 0.7 μM for niraparib and 20 μM for RI-1. Adding niraparib to TMZ increased the efficacy of the alkylating agent in both cell lines, yet with a more pronounced increment, as compared to TMZ single agent therapy, in MGMT+ than in MGMT-cells (3-way ANOVA: Niraparib p<0.0001; cell line*niraparib, p=0.0019). On the opposite, the synergistic effect of RI-1 was not significant which may result from the high variability observed in RI-1 single agent efficacy for this range of concentrations (3-way ANOVA RI-1 p=0.11). However, even at this moderate dose, adding RI-1 to TMZ and niraparib exposure impacted the efficacy of the drug combinations for TMZ doses within the range of concentrations corresponding to that measured in the plasma of patients, i.e. 0-25 μM (3-way ANOVA on AUCs restricted to 0-25 μM, RI-1 p=0.035).

**Figure 7:**
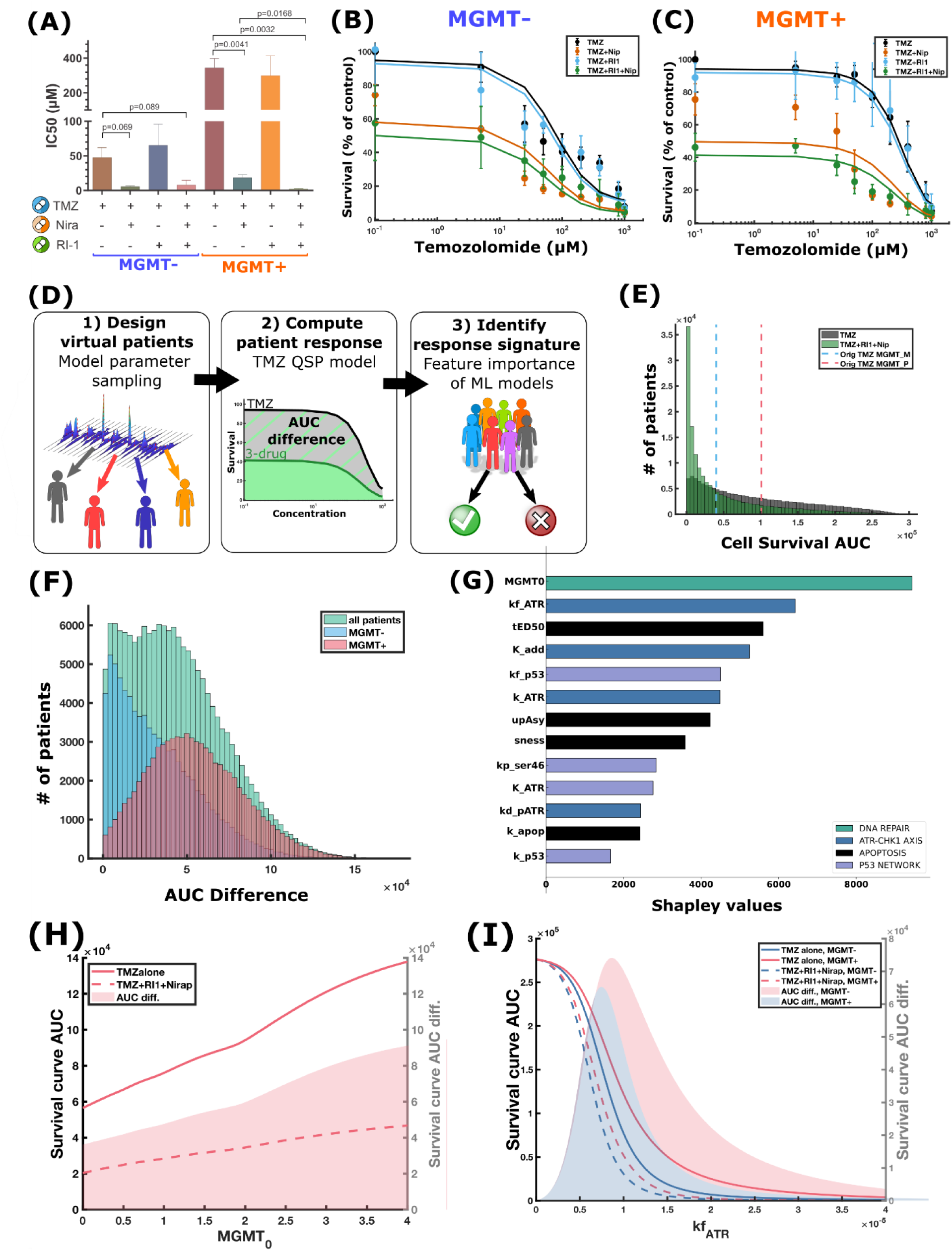
Experimental validation and functional response signature of model-derived optimal drug combinations. **(A-C)** Experimental results of the cytotoxicity of TMZ, combined or not to niraparib (BER inhibitor) or RI-1 (HR inhibitor); IC50 values (A) and dose-response curves in MGMT-(B) and MGMT+ (C) cell lines. For TMZ single agent, the solid curve represents the model best-fit after partial re-calibration. For the other conditions, model parameters were kept identical and only the levels of BER and HR inhibition were estimated from these data points. **(D)** Scheme of response signature design using virtual patients (VP) **(E)** Distributions of AUC for TMZ or the 3-drug combination across VP population. **(F)** Distributions of differences of TMZ and 3-drug AUC across MGMT- and MGMT+ VP. **(G)** Signature of response to 3-drug combination, reported as importance of model parameters to predict difference in AUC. **(H-I)** Simulations of the impact of protein levels of MGMT (MGMT_0_, (H)) and ATR (kf_ATR_, (I)) on TMZ AUC, 3-drug AUC and their difference in indicated VP populations. Other model parameters are kept identical. Surfaces correspond to AUC differences (right y-axis).

These experimental results were then used to further validate the TMZ model. As cell survival obtained in these experiments quantitatively differed from the ones used for model calibration (Fig. 3(F)), the data for TMZ single agent was used to partially recalibrate the model, re-estimating the 15 most important parameters according to the feature importance analysis (Fig. 7**(B-C)**). The three other conditions were predicted by the model by only estimating BER and HR inhibition levels corresponding to niraparib and RI-1 exposure respectively, and without modifying other model parameters. For each drug, the inhibition level was assumed to be the same for single agent or combined administration and for both cell lines. A good agreement was obtained between the data and the model, except for the combination of TMZ and niraparib in MGMT+ cells which cytotoxicity was overestimated by the model. The inhibition levels were estimated to 40% for BER and 7% for HR.

### Functional signature for benefit of the 3-drug combination over TMZ single-agent therapy

To investigate the translation of our findings to other GBM tumor models or patients, we sought to develop a response signature for the optimal 3-drug therapeutic strategy. More precisely, we determined which tumor features were associated with a benefit in terms of efficacy, of adding BER and HR inhibitors to TMZ as compared to administering the cytotoxic drug as a single agent. To investigate this, populations of MGMT+ and MGMT-virtual patients (VPs) were generated by sampling model parameters from distributions which were extended as compared to those obtained for LN229 cell lines, to account for larger patient heterogeneity (Fig. 7**(D)**, Table S5). Next, the TMZ model was used to compute the dose-response curve for each VP for either TMZ or the 3-drug combination, from which AUC values were computed. Considering all VPs, the AUC distribution was drastically skewed towards the left when adding the inhibitors to TMZ administration, indicating lower AUC values, i.e. greater efficacy of the 3-drug combination as compared to TMZ monotherapy on average over the whole VP population (Fig. 7**(E)**). The benefit of adding the targeted therapies to TMZ was quantified by the difference between the AUC of TMZ and of the 3-drug combination for each individual VP (Fig. 7**(F)**). A significant proportion of MGMT-VPs displayed small AUC differences thus suggesting a minor benefit associated to adding BER and HR inhibitors for them who may already respond to TMZ. To explore the tumor features associated with a superiority of the 3-drug combination over TMZ monotherapy, we then used machine learning (ML) to predict AUC differences from VP parameters (see Methods). ML regressors managed to predict VP AUC differences from VP parameters as R^2^ scores were equal to 0.83. The most important parameter determining the benefit of the 3-drug combination over TMZ single agent therapy was the tumor level of MGMT protein, followed by parameters involved in the ATR-CHK1 axis, p53 network and apoptosis launching (Fig. 7**(G)**). To further explain this finding, we used the TMZ model to simulate the evolution of drug AUCs for increasing values of MGMT protein level (MGMT_0_), without modifying the other parameters (Fig. 7**(H)**). As expected, the tumor resistance to TMZ and to the 3-drug combination increased with MGMT protein level as indicated by increasing AUCs. However, interestingly, the benefit of the drug combinations also increased with MGMT level thus suggesting that the patient presenting the highest overexpression of MGMT may also benefit the most from the synergy of TMZ with BER and HR inhibitors. The second most important parameter was ATR translation rate (kf_ATR_) which drives ATR intracellular concentration. In the model simulations, ‘extreme’ virtual patients with very low ATR levels were resistant to both TMZ and to the 3-drug combination because of non-functional ATR-CHK1 axis, whereas patients with very high ATR concentrations did not benefit from the addition of the targeted molecules as they were already responding to TMZ (Fig. 7**(I)**). The range of ATR concentrations resulting in a benefit of the 3-drug combination over TMZ monotherapy was wider for MGMT+ patients as compared to MGMT-VPs.

## DISCUSSION

We designed a core model of TMZ cellular PK-PD and antitumor efficacy representing the key regulatory pathways identified as either genetically altered in GBM or involved in TMZ resistance. A large effort has been made to obtain a quantitative accuracy of the model and the final model successfully recapitulated multi-type datasets for a large variety of TMZ doses and exposure durations in a sensitive and a MGMT-overexpressing resistant cell lines. After this careful model calibration, the framework was employed to design optimal therapeutic strategies enhancing TMZ efficacy in resistant cells, while preserving efficacy in initially sensitive cells. Model-derived multi-agent regimens achieving the best efficacy consisting in combining TMZ with inhibitors of BER and HR pathways, with inhibitors being administered prior to the alkylating agent. This drug combination was validated in experiments as the addition of both targeted therapies drastically reduced TMZ IC50 in both cell types, thus circumventing the initial resistance of MGMT+ cells. Using our QSP model in a machine learning pipeline, we further derive a response signature to this optimal drug combination and concluded that patients benefiting from the addition of both targeted therapies as compared to TMZ single-agent therapy must hold tumors with high MGMT concentration, functional ATR-CHK1 axis, P53 network and apoptosis pathway.

Synthetic lethality involving the administration of PARP inhibitors (PARPi) that impair BER pathways to patients with tumors bearing mutations of BRCA genes associated to a loss of function of HR repair have been successfully investigated and translated to the clinics for several cancer types including ovarian, breast and lately pancreatic cancers (*28*). However, only a subset of solid tumors bears BRCA genes mutations and alternative approaches are thus need for the other patients. For GBM, patients with tumors mutated for either BRCA1 or 2 counted for 2.4% of all TCGA patients (*29*). Recently, interest has been raised for the combination of PARPi with other targeted therapies in context not previously considered susceptible to BER inhibition (*30*). The strategy of combining both BER and HR inhibitors have been investigated in a phase I clinical trial in which ovarian cancer patients were administered PARPi and HSP90 inhibitors (*31*). This study concluded to the safety of the drug combinations and to evidence of preliminary anti-tumor efficacy. More generally, the use of PARP and other DDR inhibitors to render GBM cells more vulnerable to conventional treatments is an area of intense investigation (*32*). Our study further supports this effort as it concluded on the optimality of this drug combination, which achieved the best efficacy against MGMT-overexpressing cells, among all targeted therapies considered in this work.

The current core model includes the main proteins involved in TMZ resistance although other pathways may be relevant to add as GBM frequently involves genetic alterations of receptor tyrosine kinases for instance. As large QSP models may be hard to calibrate, an interface with bioinformatics and ML is here envisaged to prioritize the pathway to connect to the core model in a patient-specific manner (*5*). High-throughput datasets informing on plasticity or resistance mechanisms (e.g. perturbed proteomics or CRISPR-Cas9) may be analyzed through pathway enrichment (e.g. GSEA, ROntoTools(*33*)) or dimension reduction algorithm (e.g. Independent Component Analysis (*34*)) to determine the key protein networks likely to drive TMZ resistance in particular patient-derived cell lines which need to be added in the model. The most important pathways may then be connected to the core model using tools such as STRING or OMNIPATH (*35*) and the patient-specific model can be used *in silico* therapeutic optimization to identify optimal drug targets and drug combinations. Hence, the present study provides a crucial core model that will be at the heart of a wider framework aiming to derive patient-specific optimal drug combinations.

Moreover, although focused on GBM and TMZ, the approach may be applied to other malignancies as GBM is recognized as a relevant disease to represent other cancers and, as such, was the first tumor to be genetically mapped by TCGA (*36*). Indeed, several pathways represented in the model are relevant for other cancer types. The framework could serve as a basis to optimize the combinations of other cytotoxics with targeted therapies.

The limitations of this study lays in its *in vitro* nature as tumor micro-environment including blood vasculature, the blood brain barrier, and immune cells within the tumors may play a critical part in drug efficacy and may also be targeted. As in a multiscale pipeline, the core model designed here may be included as a component of a wide model describing the tumor and the drug whole-body PK. Such extension will allow to investigate other targets related to the drug brain disposition and the interactions of tumor cells with their micro-environment. In addition, the constraint of acceptable tolerability of studied drug combinations was unforced here though the minimization of drug concentrations. However, proper modeling and data may be added to the study to achieve an optimal balance between maximizing efficacy and minimizing side effects of antitumor therapies.

## MATERIALS AND METHODS

### Study design

A QSP representation of TMZ PK-PD and key regulatory protein networks involved in the drug response was developed based on existing modeling works and on the integration of multi-type datasets (Fig. 1). The model was calibrated to represent either a sensitive or a MGMT-overexpressing resistant GBM cell line. Next, numerical investigations using both models were undertaken to design optimal TMZ-based drug combinations to circumvent TMZ resistance. The model-derived optimal drug combination was validated in dedicated experiments. Next, we used machine learning regressors to derive a signature for responders of this optimal drug combination.

### Mathematical modeling and parameter estimation

TMZ model is based on ordinary differential equations (ODEs) representing chemical reactions by the mass action law for passive reactions and Hill kinetics for cooperative binding between several elements (Supplementary Information for full description). The model was solved using the ode23tb solver of Matlab (R2021b, MathWorks). Model calibration was performed in a two-step process in which the first step consisted in estimating subsets of parameters when data availability permitted it (Supplementary Figures and Table S2-S3). For the second step, model parameters were estimated using a modified least square approach in which the L2 goodness-of-fit is supplemented with semi-quantitative constraints as follows:

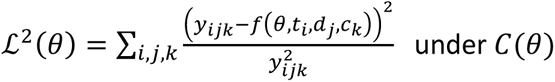

where *y*_*ijk*_ is the data at time *t*_*i*_, at the dose *d*_*j*_ of TMZ in the LN229 clone *c*_*k*_ ∈ {*MGMT*−, *MGMT* +}, and *f*(*θ*, *t*_*i*_, *d*_*j*_, *c*_*k*_) is the value computed by the model at time *t*_*i*_, at the dose *d*_*j*_ of TMZ related to the LN229 clone *c*_*k*_. To increase the parameter identifiability, several constraints gathered in *C*(*θ*) were imposed to ensure biologically sound parameter values (Supplementary Information). Numerical minimization was performed by the Covariance Matrix Evolutionary Strategy (CMA-ES) (*37*). Initial guess for parameter values were fixed according to biological and physical knowledge found in the literature when available. Next, the algorithm was run in an iterative process in which initial guesses and search intervals were updated at each loop according to the best parameter set obtained in the preceding run.

### Experimental Data used for parameter estimation

The model was calibrated using several published time- and TMZ dose-dependent datasets (Supplementary Information). For the final step of parameter estimation, the data was available in both LN229 wild type, not expressing MGMT and here denoted with MGMT-, and a LN229 cell line stably modified to express MGMT, here denoted with MGMT+. Mainly four studies were integrated, described here. In Stratenwerth et al. (*27*), O6-methyl-deoxyguanosine adducts were quantified for different TMZ doses after 3h of exposure (Fig. 3 (A)) and in a longitudinal manner during TMZ exposure (50 μM, unshown). In He et al. (*26*) were published: dose-dependent *γ*H2AX foci assay after 72 h of TMZ exposure (Fig. 3(B)); TMZ dose-dependent WB analysis of P53 and p-P53-Ser46 expression after 24 h and 72 h of exposure (Fig. 3(C,D)); TMZ dose-dependent apoptotic cell percentage after 72 h, 96 h, 120 h and 144 h of exposure (Fig. 3(E)). In Jackson et al. (*24*), several datasets were available including: time-dependent western blot (WB) analysis of MGMT, CHK1 and pCHK1 expression levels after exposure to 12.5 μM or 100 μM of TMZ (Fig. 4 (C,I,J)), or to various doses of ATR inhibitor VX970 (Fig. S5); cell survival after six days of exposure to TMZ at various concentrations (Fig. 3 (F)). In Aasland et al. 2020 (*25*), we could retrieve: time-dependent WB analysis of pCHK1, CDC25, P53 and P53ser46 expression after exposure to 100 µM of TMZ (Fig. 4 (J, K, M, N)).

### Vine copula to estimate parameter multi-variate distribution

A vine copula was used to estimate the multivariable distribution of the 25 parameters estimated in the final step of model calibration. Parameter sets yielding the 1% best cost function values as computed by the CMAES algorithm were used to calibrate the copula model (i.e. 90111 sets of 25 parameters). The vine copula model definition and calibration were performed using the R library “rvinecopulib”. First, multiple models for univariate marginals were tested and the best one was selected using the AIC criteria for each parameter. Then, the parameter distributions lying in the unit hypercube are used to estimate the vine copula model using parametric families. The marginal distributions and estimated parameters are reported in Table S4.

### Parameter Importance and gradient boosting models

Parameter importance was determined taking as output the AUC of cell survival curve after 6 days of TMZ exposure. A selection of 28 out of the 69 model parameters were tested excluding PK parameters not influencing the model output (i.e. p_A_ and p_A2_), all Hill exponents, phenomenological parameters of the S-phase marker and including only one parameter acting on each protein total amount (i.e. including only transcription rates and not degradation rates). Three of them were cell-line specific, meaning that their value differed between MGMT- and MGMT+ cells. Multiple parameter sets were generated using the vine copula model for 18 parameters and assuming log-norm distributions with means equal to the best fit estimates and coefficients of variations of 20% for others (R library “rvinecopulib”, Table S4). The number of parameter sets was estimated using the meaning curve strategy and set to 80,000, 80% of them being used for training and 20% for validation. Next, these parameters were input to the QSP TMZ model to compute the corresponding AUC of cell survival curve 6 days after TMZ exposure. This step was performed using Matlab.

Gradient boosting models were fitted to predict TMZ AUCs from parameter values (“scikit-learn” library in Python). The hyperparameter were tuned by performing a 10-fold cross validation using grid search (*GridSearchCV* function) and based on 64000 parameters sets for each MGMT+ or MGMT-condition. Four hyperparameters were optimized: the *loss* function (tested value [’squared_error’, ‘absolute_error’, ‘quantile’, ‘huber’], the number of trees (*n_estimator*, searched in [10, 100, 500]), the maximum depth of a single tree (*max_depth*, searched in [2, 7, 10, 13, None]), and the maximum features of a single tree (*max_features*, set to either None or to the square root of the total number of features). MGMT- and MGMT+ AUCs were modeled separately and the optimal hyperparameter values obtained were the same and equal to *loss*: ‘squared_error’, *n_estimator*: 500, *max_depth*: 7, *max_features*: None. The prediction R^2^ scores computed on the 16000 parameter sets kept for validation were equal to 0.95 and 0.86, for MGMT+ and MGMT-respectively. Finally, feature importance of the calibrated gradient boosting model was assessed using shapley values (*Explainer* function of *shap* package).

### Therapeutic Optimization

The calibrated QSP TMZ model was used to investigate drug combinations *in silico*. To simulate the activity of a targeted inhibitor, the model parameter corresponding to the target was decreased by some percentage, reflecting the compound dose. For instance, exposure to an inhibitor of the BER pathway was represented as a decrease in the k_BER_ parameter. Parameter values were modified from the initial time and remained constant over the whole TMZ exposure thus assuming that the inhibition had reached steady state at time 0 and remained active over time.

### Virtual Patient Population and gradient boosting models

A virtual population of patients was generated as follows. A patient was defined by a set of 29 parameters sampled from log normal distribution with extended standard deviations as compared to LN229 cell lines estimates, to account for inter-patient variability (R library “rvinecopulib”, Table S5). Next, for a given VP, i.e. a given parameter set, the TMZ model was used to compute the AUC of cell survival curve after 6 days of exposure to TMZ given as a single agent or combined to BER and HR inhibitors using Matlab. Learning curve analysis concluded that a population of 160000 VPs (80000 for each MGMT+ or MGMT-condition) was needed. Eighty percent of the virtual population was used as a training dataset to calibrate gradient boosting regressors predicting the AUCs from parameter values, and the remaining 20% virtual patients were used as a validation cohort.

We used a gradient boosting model to predict AUC of different treatments for all patients using the *GradientBoostingRegressor* function from the “scikit-learn” library in Python. To tune hyperparameters, we performed a 10-fold cross validation using grid search (*GridSearchCV* function). Three hyperparameters were optimized: the number of trees (*n_estimator*, searched in [10, 100, 500]), the maximum depth of a single tree, (*max_depth*, searched in [None, 2, 8,10, 12]), and the maximum features of a single tree (*max_features*, set to either None or to the square root of the total number of features). The resulting optimal hyperparameter values were *n_estimator*: 500, *max_depth*: 12, *max_features*: None. The prediction R^2^ score computed on the validation VP cohort was equal to 0.83. Once the gradient boosting regressors were calibrated, we performed a Feature Importance (FI) analysis to identify parameters which impacted the most on the model output, based on shapley values (*Explainer* function of *shap* package).

### Experimental validation of optimal drug combinations

LN229 MGMT- and MGMT+ cell lines were kindly donated by Dr. Bernd Kaina (Institut für Toxikologie, Universitätsmedizin, Mainz, Germany). TMZ (#T2577) was purchased from Sigma. Niraparib (#TA-T3231) and RI1 (#TA-T2276) were purchased from Euromedex. LN229 cells were seeded in 96-well plates (3000 cells per well) and maintained in DMEM 10% fetal bovine serum at 37 °C, 5% CO_2_. Twenty-four hours after cell seeding, drugs were added at indicated concentrations, and cells were incubated for 96 hours. At the end of the incubation period, cell viability was measured using the wst-1 assay (CELLPRO-RO, Sigma) according to the manufacturer’s instructions. Results are reported as percentages of treated conditions over control conditions corresponding to cells cultured in regular culture medium.

### Statistical tests

AUC and IC50 of cell viability data were computed with GraphPad. IC50s were determined using the non-linear regression ECanything equation. Three-way ANOVA and paired t-test were performed using GraphPad.

## Acknowledgments

We acknowledge Prof. Dr. Bernd Kaina (Institut für Toxikologie, Universitätsmedizin, Mainz, Germany) for providing the LN229 cell lines and Dr Niklas Hartung (Institut für Mathematik, Potsdam,DE) for fruitful discussion and assistance regarding the vine copula approach.

## Author contributions

Conceptualization: SC, MV, AI, AB.

Methodology: SC, MV, HM, CC, AI, AB.

Investigation: SC, MV, HM, TD, CC, AI, AB.

Visualization: SC, MV, TD, AB.

Funding acquisition: MV, AI, AB.

Project administration: MV, AB.

Supervision: MV, AI, AB.

Writing – original draft: SC, MV, TD, AI, AB.

Writing – review & editing: SC, MV, TD, AI, AB

## Data and materials availability

All datasets, Matlab and R codes are available at https://github.com/SyspharmaCurie/Corridore-et-al.-Codes.

