## Supplementary Information for "Circumventing glioblastoma resistance to temozolomide through optimal drug combinations designed by quantitative systems pharmacology and machine learning"

#### Outline


##### 1. TMZ cellular PK-PD model description and Step 1 of parameter estimation

1.1 TMZ pharmacokinetics

1.2 DNA adducts formation

1.3 MGMT repair mechanisms

1.4 Mismatch Repair

1.5 Base Excision Repair

1.6 DNA double stranded breaks dynamics

1.7 DNA damage response and cell cycle arrest

1.8 The cell cycle module

1.9 P53 – MDM2 signaling

1.10 The cell population module

##### 2. Steady State Assumptions

##### 3. Step 2 of Model Parameter Estimation

3.1 Parameter estimation strategy

3.2 Biological constraints applied to parameter estimation

### 1 TMZ cellular PK-PD model description and preliminary parameter estimation

In this section, we describe in detail the TMZ cellular PK-PD model presented in the main text, together with *step 1* of parameter estimation. Model variables and parameters are defined in Table S1 and S2 respectively.

#### 1.1 TMZ pharmacokinetics

The previously-published model of TMZ pharmacokinetics (PK) *(1)* describes TMZ cell membrane transport and metabolism in the intracellular and extracellular medium. TMZ is activated into its metabolite 5-(3-methyltriazen-1-yl) imidazole-4-carboxamide (MTIC) and MTIC is subsequent degraded into 4-amino-5-imidazole-carboxamide (AIC)—an inactive metabolite—and a methyldiazonium cation, the DNA-methylating species, those two reactions being pH-dependent. Because TMZ is highly lipophilic and constitutes a poor substrate of ABC transporters, transport of the parent drug and AIC between the extracellular and intracellular compartments were modeled as passive diffusion. As MTIC displays limited ability to cross cell membranes and as the methyldiazonium cation is a highly reactive species, its transport between both spatial compartments were not considered. Here are the equations for TMZ PK:

$$\frac{dT_{out}}{dt} = -\frac{p_T * T_{out} - p_{T2} * T_{in}}{V_{out}} - k_T^{out} T_{out} \,,$$

$$\frac{dT_{in}}{dt} = \frac{p_T * T_{out} - p_{T2} * T_{in}}{V_{in}} - k_T^{in} T_{in} \,,$$

$$\frac{dM_{out}}{dt} = k_T^{out} T_{out} - k_M^{out} M_{out},$$

$$\frac{dM_{in}}{dt} = k_T^{in} T_{in} - k_M^{in} M_{in} \,,$$

$$\frac{dA_{out}}{dt} = -\frac{p_A * A_{out} - p_{A2} * A_{in}}{V_{out}} + k_M^{out} M_{out} \,,$$

$$\frac{dA_{in}}{dt} = \frac{p_A * A_{out} - p_{A2} * A_{in}}{V_{in}} + k_M^{in} M_{in} \,,$$

$$\frac{dC_{in}}{dt} = k_M^{in} M_{in} - \left(\frac{1}{2} k_{addO} + k_{addN}\right) DNA\ C_{in} - k_{cat} C_{in} \,,$$

where $V_{in}$ and $V_{out}$ are the intra- and extra-cellular volumes. Equations on $A_{out}$ and $A_{in}$ were discarded as they do not impact the rest of the model. All PK parameters were inferred from the previous study based on U87 glioma cell culture PK data *(1)*.

#### 1.2 DNA adduct formation

Concerning TMZ pharmacodynamics (PD), the methyldiazonium cation $C$ is the sole species able to form DNA adducts. $C$ interacts with guanine (G) and adenine (A) bases at N7-G, N3-G, N3-A and O6-G, to create N7-methylguanine (N7-meG), N3-methylguanine (N3-meG), N3-methyladenine (N3-meA) and O6-methylguanine (O6-meG) DNA adducts. Since N7-meG, N3-meA and N3-meG adducts are all repaired by the same repair system (i.e. Base Excision Repair, here BER), we only modeled the sum of these three species, denoted as N37-meAG. N37-meAG and O6-meG are created at respective rates $k_{addN}$ and $k_{addO}$.

In the absence of drug, regular intracellular processes may create basal DNA damage and DSBs which were accounted for in the model. Experimental data showed that DSBs were present at very low TMZ doses in similar amount in LN229 MGMT+ and MGMT- cells suggesting that this basal damage was independent of O6-meG adduct dynamics (Figure 2B). For this reason, and for the sake of simplicity, to avoid adding another variable/equation, we lumped DNA damage produced independently of TMZ with N37-meAG adduct induced by the drug. We assumed that these TMZ-independent damage were formed at a constant rate $k_{addEx}$ added in N37-meAG equation and were repaired by BER. Here are the equations governing their dynamics:

$$\frac{dO6mG}{dt} = \frac{1}{2} k_{addO} DNA\, C_{in} - k_{MGMT} O6mG\, MGMT - k_{GT}\, O6mG \left(1 + k_{CCNA} \frac{CCNA^{n_{cc}}}{CCNA^{n_{cc}} + {K_{cc}}^{n_{cc}}}\right),$$

$$\frac{dN37mAG}{dt} = k_{addEX} + k_{addN} DNAC_{in} - k_{BER}\, N37mAG - k_{DSBN}\, N37mAG \left(1 + k_{CCNA} \frac{CCNA^{n_{cc}}}{CCNA^{n_{cc}} + {K_{cc}}^{n_{cc}}}\right).$$

As previously computed *(1)*, *DNA* base pairs are in large excess compared to the intracellular methylating cation concentration so that the concentration of A and G bases could be considered as constant in the model. DNA base pairs concentration (either A-T or G-C) was estimated to 5500 µM in a cell of 1pL *(1)*. For the formation of O6meG adducts, only G-C pairs are targeted so that a factor of one half was added.

#### 1.3 MGMT repair mechanisms

O6-meG adducts are specifically repaired by MGMT which gains the methyl group initially attached to the DNA base in a stoichiometric manner as one molecule of MGMT is consumed to repair one adduct. This reaction is known to follow the law of mass action *(2–4)*, with a reaction rate that we denoted $k_{MGMT}$. We also considered that MGMT protein is constantly produced –

constant rate $k_f^{MGMT}$ – and degraded –constant rate $k_d^{MGMT}$. Here is the equation for MGMT protein concentration:

$$\frac{dMGMT}{dt} = k_f^{MGMT} - k_d^{MGMT} MGMT - k_{MGMT}\, O6mG\; MGMT\ .$$

This equation was not considered for MGMT- cell in which MGMT levels were known to be zero. MGMT protein half-life was evaluated in T98G glioma cells to 63h *(6)* which allowed us to compute the degradation parameter: $k_d^{MGMT} = \frac{\ln(2)}{63} = 0.011\ h^{-1}$. This value was in agreement with a study in NIH-3T3 mouse fibroblasts *(5)* in which they estimated MGMT half-life to 34.36h which would mean $k_d^{MGMT} = 0.02\ h^{-1}$. On the opposite, authors from *(7)* claimed that they could retrieve MGMT half-life from the study in *(8)* but mistakenly used measurements performed in the presence of methylating agents which artificially decreased MGMT half-life to approximately 2h, leading to an overestimation of MGMT protein degradation constant rate. Next, $k_{MGMT}$ was estimated in step 2 of parameter estimation, together with MGMT initial concentration ($MGMT_0$) and the production rate $k_f^{MGMT}$ was computed using steady-state assumptions as explained here below.

##### 1.4 Mismatch Repair

The presence of O6-meG adducts on the DNA triggers the mismatch repair (MMR) machinery in a dysregulated manner. Indeed, the O6-meG:C base pairs are mispaired during replication due to the presence of the methyl group, the C base is replaced by a T base and O6-meG:T are created *(9)*. This mispair is detected by the MMR system which excises the T base but immediately re-inserts one in the DNA due to the mispair properties of O6-meGs. Those futile loops result in persistent Single Stranded Breaks (*SSB*) on the DNA. Then, replication forks collide with SSBs which creates Double Stranded Breaks (*DSB*). In the model, O6-meG are transformed into O6-meG:T (*GT*) at rate $k_{GT}$ when the cell is in S-phase (see cell cycle module below). Then, O6-meG:T are transformed into single stranded breaks (*SSB*) by the MMR system at the rate $k_{MMR}$. Thus, we do not detail the MMR steps of the futile loops leading to the persistent SSB assuming that they occur very rapidly compared to the other reactions. During the S-phase, SSB are transformed into DSB at the rate $k_{DSBO}$. Here are the corresponding equations:

$$\frac{dGT}{dt} = k_{GT}\ O6mG\left(1 + k_{CCNA}\frac{CCNA^{n_{cc}}}{CCNA^{n_{cc}} + {K_{cc}}^{n_{cc}}}\right) - \ k_{MMR} GT\ ,$$

$$\frac{dSSB}{dt} = \ k_{MMR} GT \quad - k_{DSBO}\ SSB\left(1 + k_{CCNA}\frac{CCNA^{n_{cc}}}{CCNA^{n_{cc}} + {K_{cc}}^{n_{cc}}}\right),$$

Experimental data from *(10)* used in Step 2 of parameter estimation only inform on the sum of O6me, GT and SSB, and adding up the equations of these three variables provides an equation where the parameters $k_{GT}$ and $k_{MMR}$ are no more present, thus making them practically unidentifiable. The parameter $k_{GT}$ was arbitrarily set to one as no information could be found in the literature about it. For $k_{MMR}$ estimation, we used the work of Geng and colleagues *(11)* who assessed MMR in vitro kinetics by incubating two different circular plasmids containing a mismatch G:T, pSCW01 GT DNA and pSCW02 GT DNA, with MMR proteins and monitoring their disappearance. Seventy-five fmol of each circular plasmid were placed in 40μL of assay medium and repair proteins were added at time t=0. After 7 min of incubation, the remaining fraction of mismatch DNA were equal to 57% and 45%, for pSCW01 GT and pSCW02 GT, respectively, which led to a mean ± SD of 52 ± 8.5%. Assuming an exponential decay in GT mispair concentration, with rate $k_{MMR}$ and plugging this data into the equation, led to an estimation of $k_{MMR}$ = 5.8 ± 1.04 $h^{-1}$.

#### 1.5 Base Excision Repair

In the model, the Base Excision Repair (BER) system repairs N37-meAG adducts and basal DNA damage occurring in the absence of drug *(9)*. In the event of deficient BER system, N37-meAG adducts may induce DNA lesions in the form of DSBs. However, the BER system is usually efficient and the main cytotoxicity associated to TMZ is thought to be the consequence of O6-meG adducts *(12)*. N37-meAG adduct repair by the BER system is represented by a linear degradation of rate $k_{BER}$ , thus lumping the entire pathway into a single parameter for the sake of simplicity. $k_{BER}$ was estimated from in vitro kinetics experiments (Figure 2A of *(13)*). The recognition of adducts by the BER system occurred in vitro at a rate of 0.6 nM.$s^{-1}$ with a large concentration of DNA adducts of 1 $\mu$M. Hence, $k_{BER}$ was set to 0.6 $s^{-1}$ or 2.16 $h^{-1}$. Remaining adducts may create DSBs through collisions with replication forks during the cell S-phase at the rate $k_{DSBN}$ .

#### 1.6 DNA double stranded breaks dynamics

Double Stranded Breaks (DSB) originating from upstream DNA damage are repaired by Homologous Recombination at the rate $k_{HR}$. Here is the equation governing their dynamics:

$$\frac{dDSB}{dt} = (k_{DSBO}SSB + k_{DSBN}N37mAG)\left(1 + k_{CCNA}\frac{CCNA^{n_{cc}}}{CCNA^{n_{cc}} + {K_{cc}}^{n_{cc}}}\right) - k_{HR}DSB\ ,$$

We utilized the time-resolved experimental data of H2AX foci measurement from Figure 2 of *(14)* to estimate the rate of DSB repair and found $k_{HR} = 0.166h^{-1}$. It was consistent with the value estimated from lung cancer cell culture from Figure1C-D of *(15)* which gave $k_{HR} = 0.175h^{-1}$.

#### 1.7 DNA damage response and cell cycle arrest

The sensing of TMZ-induced DNA damage seems to be primarily driven by ATR as ATM played only a minor role in early hours after drug exposure *(16–18)*. ATR exists as unphosphorylated proteins in the absence of DNA damage and is activated (p*ATR*) by the detection of replication fork stalling leading to DSBs through successive protein-protein interactions that we do not explicitly model here *(17, 19–21)*.

In the model, unphosphorylated ATR protein is constantly produced at constant rate $k_f^{ATR}$and degraded, at constant rate $k_d^{ATR}$. ATR activation occurs only once a minimum amount of DSB is reached, such reaction being represented by a Hill function of DSB amounts with parameters $k_{ATR}$, $K_{add}$ and $n_{add}$. Moreover, pATR is supposed to be continuously degraded at rate $k_d^{pATR}$. Once activated, ATR kinases phosphorylate CHK1 proteins into pCHK1 *(16)*. Similar to ATR, we also considered a continuous production and degradation of CHK1 with respective constant rates $k_f^{CHK1}$ and $k_d^{CHK1}$, and continuous degradation of pCHK1 at rate $k_d^{pCHK1}$. pCHK1 enhances the degradation of CDC25 phosphatases, at constant rate $k_{CDC25}$, leading to cell cycle arrest *(16)*. Indeed, TMZ is known to induce cell cycle arrest during the S-phase and at the G2/M checkpoint. In the model, we represented CDC25 production and degradation with constant rates, $k_f^{CDC25}$and $k_d^{CDC25}$. Here are the corresponding equations:

$$\frac{dATR}{dt} = k_f^{ATR} - k_{ATR}\,\frac{DSB^{n_{add}}}{DSB^{n_{add}} + {K_{add}}^{n_{add}}}\,ATR - k_d^{ATR} ATR\,,$$

$$\frac{dp\mathrm{ATR}}{dt} = k_{ATR}\,\frac{DSB^{n_{add}}}{DSB^{n_{add}} + {K_{add}}^{n_{add}}} ATR - k_d^{pATR} pATR\,,$$

$$\frac{dCHK1}{dt} = k_f^{CHK1} - k_{CHK1}\,CHK1\,pATR - k_d^{CHK1} CHK1\,,$$

$$\frac{dpCHK1}{dt} = k_{Chk1}\,CHK1\,pATR - k_d^{pCHK1} pCHK1\,,$$

$$\frac{d\,CDC25}{dt} = k_f^{CDC25} - k_{CDC25}\,pCHK1\,CDC25 - k_d^{CDC25} CDC25\,,$$

The parameters $k_f^{ATR}$, $k_d^{ATR}$, $k_f^{Chk1}$, $k_d^{CHK1}$, $k_f^{CDC25}$ and $k_d^{CDC25}$ were directly taken from a quantitative proteomics study in NIH-3T3 mouse fibroblasts *(5)*.

#### 1.8 The cell cycle module

In the TMZ PK-PD model, the only needed cell cycle input lays in the information of the cell being or not in S-phase. Therefore, the cell cycle module could be reduced to a S-phase marker. We chose to base this marker on the cyclin A (CCNA) protein level as it transiently increases during the S phase of cycling cells. This can also be seen in a detailed mathematical model of the cell cycle that shows the particular shape of cyclin A protein amount along the cell cycle *(22)*. Thus, we modeled cyclin A level as the following cosine function:

$$CCNA(t) = \left(M_{cc} + A_{cc}cos\left(\frac{2\pi}{T_{cc}}(t-\varphi)\right)\right)\frac{CDC25^{n_{cc}}}{CDC25^{n_{cc}} + K_{CDC25}^{n_{cc}}},$$

where t is the time, $M_{cc}$ is CCNA average level, here set to 1, $A_{cc}$ is its amplitude, with value set to 1, $\varphi$ is its phase, and $T_{cc}$ is the period set to 24h as the cell cycle is usually entrained by the circadian clock. The S-phase marker was obtained by computing the following Hill function of CCNA:

$$S = k_{CCNA}\frac{CCNA^{n_{cc}}}{CCNA^{n_{cc}} + {K_{cc}}^{n_{cc}}}.$$

Parameters $K_{cc}$ and $n_{cc}$ were set so that S was close to 1 in S-phase and close to zero elsewhere. For a cell cycle of period 24h, we considered that the S-phase last 6h, that is 1/4 of the total cell cycle, and occurred when CCNA was at its maximum values (i.e. in [$\varphi$ $-3h$, $\varphi$ + 3h] modulo $24h$). The phase of CCNA $\varphi$ was set to 9h so that, at t=0, the cell was in G1, and the first S-phase occur between 6h and 12h of TMZ exposure.

The cell cycle arrest was represented by a Hill function of CDC25 protein levels which rapidly set CCNA levels to zero in the absence of the phosphatases. The activation threshold and the Hill coefficient are denoted respectively by $K_{CDC25}$ and $n_{cc}$.

##### 1.9 P53 – MDM2 signaling

The signaling pathway of the P53 tumor suppressor protein was represented by a modified version of the model of Sturrock et al. 2011 *(23)*. Briefly, in standard no-stress conditions (no TMZ injection), P53 is present in the cell at low level. In response to TMZ-induced cellular stress, P53 levels are increased through an enhancement of its translation. Moreover, TMZ causes the phosphorylation of P53 at serine 46, producing P53-Ser46 species which is a promoter of pro-apoptotic genes *(21)*. P53 and P53-Ser46 then promote the transcription of MDM2. In turn, MDM2 negatively regulate P53 expression which creates a negative feedback loop.

In the model, the P53-MDM2 system is represented by four ODEs. In the ODE describing P53, P53 is assumed to be continuously produced at a constant rate $k_t^{P53}$. The translation term is enhanced by a Hill function of phosphorylated ATR (activation threshold and Hill coefficient denoted respectively $K_{ATR}$ and $n_{ATR}$), reflecting TMZ-mediated P53 level increase. The P53 degradation is composed by a natural degradation term of rate $k_d^{P53}$, and by a degradation term dependent on the amount of MDM2, in the form of a Hill function, with degradation rate $k_d^{P53MDM2}$, activation threshold $K_{MDM2}$ and Hill coefficient $n_{MDM2}$. The phosphorylation of P53 follows the law of mass action in P53 and ATR*, with a constant phosphorylation rate $k_p^{ser46}$. The

P53-Ser46 degradation occurs at constant rate $k_d^{ser46}$. The ODE modelling MDM2 mRNA has a production term with basal rate $k_t^{MDM2}$, which is enhanced by a Hill function dependent on the amount of P53 and P53-Ser46, reflecting the activity of P53 as a transcription factor, and finally a natural degradation term of rate $k_{td}^{MDM2}$. The activation threshold and Hill coefficient are denoted respectively $K_{P53}$ and $n_{P53}$. Finally, the ODE for MDM2 protein is composed of a translation term dependent on the amount of MDM2 mRNA, with constant rate $k_f^{MDM2}$, and a natural degradation term, with rate $k_d^{MDM2}$.

$$\frac{dP53}{dt} = k_t^{P53}\left(1 + k_{P53}\frac{pATR^{n_{ATR}}}{pATR^{n_{ATR}} + K_{ATR}{}^{n_{ATR}}}\right) - \left(k_d^{P53} + k_d^{P53MDM2}\frac{MDM2^{n_{MDM2}}}{MDM2^{n_{MDM2}} + K_{MDM2}{}^{n_{MDM2}}}\right)P53 - k_p^{ser46}P53\, pATR\,,$$

$$\frac{dP53ser46}{dt} = k_p^{ser46}P53\, pATR - k_d^{ser46}P53ser46\,,$$

$$\frac{dmRNA_{MDM2}}{dt} = k_t^{MDM2} + k_t^{MDM2P53}\frac{(P53 + P53ser46)^{n_{P53}}}{(P53 + P53ser46)^{n_{P53}} + K_{P53}{}^{n_{P53}}} - k_{td}^{MDM2}\, mRNA_{MDM2}\,,$$

$$\frac{dMDM2}{dt} = k_f^{MDM2}mRNA_{MDM2} - k_d^{MDM2}MDM2\,.$$

##### 1.10 The cell population module

The module comprises an ODE describing the dynamics of the number of cells over time during TMZ exposure. It is composed only of a death term, with rate $k_{apop}$, the cell proliferation being neglected as TMZ is known to stop the cell cycle.

According to *(24)*, the area under the curve of P53 protein amounts measured in single cells was highly correlated with cell death. Indeed, apoptosis was triggered once this quantity reaches a certain threshold and this threshold increased over time. Hence, the death term takes as input the cumulative amount of P53 and P53-Ser46 produced by a given TMZ dose as computed by the PK-PD model. To reproduce biological findings, we designed a Hill function of P53 AUC values, $f(t)$, which regulates the cell population decrease. Moreover, the threshold corresponding to apoptosis launching, rather than being constant, increases in time as observed in single cell experiments. The activation threshold is given by a time-dependent logistic function $K(t)$ with parameters $upAsy$ as upper asymptote, $sness$ as the stiffness and $t_{ED50}$ as the time when the function reaches 50% of its maximum value. Here are the corresponding equations:

$$\frac{dN}{dt} = -\, k_{apop}f(t)N,$$

$$f(t) = \frac{\left(\int_{t_0}^{t} P53ser46dt\right)^{n_{intP53}}}{\left(\int_{t_0}^{t} P53ser46dt\right)^{n_{intP53}} + K(t)^{n_{intP53}}},$$

$$K(t) = upAsy\left(1 - \frac{1}{1 + e^{sness(t - t_{ED50})}}\right).$$

#### 2. Steady state assumption

We assume that, at the initial time, just before TMZ administration, the system is at steady state. Thus, we computed the system steady state when $TMZ_{out}(t=0) = 0\ \mu M$, by setting all equations to zero. The steady state values of state variables are denoted with $(ss)$ superscript. The relationships exposed below were used to compute the parameters and initial conditions indicated in bold, in order to decrease the numbers of degrees of freedom in the parameter estimation steps.

**$TMZ\ PK$ equation at steady-state:**

$$T_{out}^{(ss)} = T_{in}^{(ss)} = M_{out}^{(ss)} = M_{in}^{(ss)} = C_{in}^{(ss)} = 0.$$

**$06meG$ equation at steady-state:**

$$06meG^{(ss)} = 0.$$

**$MGMT$ equation at steady-state:**

$$\boldsymbol{k_f^{MGMT}} = \frac{k_d^{MGMT}\ MGMT^{(ss)}}{mRNA_{MGMT}{}^{(ss)}}.$$

**$GT$ equation at steady-state:**

$$GT^{(ss)} = 0.$$

**$SSB$ equation at steady-state:**

$$SSB^{(ss)} = 0.$$

**$N37meAG$ and DSB equation at steady-state:**

$$\boldsymbol{N37mAG^{(ss)}} = \frac{k_{addEx}}{k_{BER} + k_{DSBN}\left(1 + k_{CyA}\frac{CCNA^{(ss)^{n_{cc}}}}{CCNA^{(ss)^{n_{cc}}} + K_{cc}{}^{n_{cc}}}\right)},$$

$$\boldsymbol{DSB}^{(ss)} = \frac{N37mAG^{(ss)}\, k_{DSBN} \left(1 + k_{CyA} \frac{CCNA^{(ss)^{n_{cc}}}}{CCNA^{(ss)^{n_{cc}}} + K_{cc}{}^{n_{cc}}}\right)}{k_{HR}},$$

where $CCNA^{(ss)}$ was computed through the Cyclin A equation at the initial time, thus switching off the cell cycle oscillating forcing function for the sake of simplicity.

$k_{addEx}$ was estimated using the equations for $N37mAG^{(ss)}$ and $DSB^{(ss)}$ together with available data stating that $\mathrm{DSB}^{(ss)}(72\mathrm{h}) = 1.165\mathrm{e}^{-5}$ μM in the absence of TMZ *(21)*. From $N37mAG^{(ss)}$ equation, it follows:

$$\boldsymbol{k_{addEx}} = \left[k_{BER} + k_{DSBN} \left(1 + k_{CCNA} \frac{CCNA^{(ss)}(72h)^{n_{cc}}}{CCNA^{(ss)}(72h)^{n_{cc}} + K_{cc}{}^{n_{cc}}}\right)\right] N37mAG^{(ss)}(72h)\,,$$

where $CCNA^{(ss)}(72h)$ corresponds to the concentration of Cyclin A at time t=72h, in the absence of TMZ, and where $N37mAG^{(ss)}(72h)$ was computed using the $DSB^{(ss)}$ equation as:

$$\boldsymbol{N37mAG}^{(ss)}(72h) = \frac{k_{HR} DSB^{(ss)}(72h)}{k_{DSBN} \left(1 + k_{CyA} \frac{CCNA^{(ss)}(72h)^{n_{cc}}}{CCNA^{(ss)}(72h)^{n_{cc}} + K_{cc}{}^{n_{cc}}}\right)},$$

**$\boldsymbol{ATR}$ equation steady-state:**

$$\boldsymbol{ATR}^{(ss)} = \frac{k_f^{ATR}}{k_{ATR} \frac{DSB^{(ss)^{n_{add}}}}{DSB^{(ss)^{n_{add}}} + K_{add}^{n_{add}}} + k_d^{ATR}},$$

with $\mathrm{k_f^{ATR}} = \mathrm{k_f^{ATR,MGMT+}}$ for LN229 MGMT+ cells and $\mathrm{k_f^{ATR}} = \mathrm{k_f^{ATR,MGMT-}}$ for MGMT- cells.

**$\boldsymbol{pATR}$ equation steady-state:**

$$\boldsymbol{pATR}^{(ss)} = \frac{k_{ATR} \frac{DSB^{(ss)^{n_{add}}}}{DSB^{(ss)^{n_{add}}} + K_{add}^{n_{add}}} ATR^{(ss)}}{k_d^{pATR}},$$

with different $ATR^{(ss)}$ for MGMT- and MGMT+ cell lines, computed with the equation above.

**$CHK1$ equation steady-state:**

$$\boldsymbol{CHK1}^{(ss)} = \frac{k_f^{CHK1}}{k_{CHK1}\, pATR^{(ss)} + k_d^{CHK1}},$$

with $\mathrm{k_f^{CHK1}} = \mathrm{k_f^{CHK1,MGMT+}}$ for LN229 MGMT+ cells and $\mathrm{k_f^{CHK1}} = \mathrm{k_f^{CHK1,MGMT-}}$ for MGMT- cells.

**$pCHK1$ equation steady-state:**

$$\boldsymbol{pCHK1}^{(ss)} = \frac{k_{CHK1}\, CHK1^{(ss)}\, pATR^{(ss)}}{k_d^{pCHK1}},$$

with different $\mathrm{CHK1^{(ss)}}$ for MGMT- and MGMT+ cell lines, computed with the equation above.

**$CDC25$ equation at steady-state:**

$$\boldsymbol{k_f^{CDC25}} = k_{CDC25} pCHK1^{(ss)}\, CDC25^{(ss)} + k_d^{CDC25}\, CDC25^{(ss)}.$$

Since $\mathrm{k_f^{CDC25}}$ depends on $\mathrm{pCHK1^{(ss)}}$, we have set different $\mathrm{k_f^{CDC25}}$ values for MGMT- and MGMT+ cells.

**$P53$ equation at steady-state:**

$$\boldsymbol{P53}^{(ss)} = \frac{k_f^{P53}\left(1 + k_{P53}\dfrac{{pATR^{(ss)}}^{n_{ATR}}}{{pATR^{(ss)}}^{n_{ATR}} + {K_{ATR}}^{n_{ATR}}}\right)}{\left(k_d^{P53} + k_d^{P53MDM2}\dfrac{{MDM2^{(ss)}}^{n_{MDM2}}}{{MDM2^{(ss)}}^{n_{MDM2}} + {K_{MDM2}}^{n_{MDM2}}} + k_p^{ser46}\, pATR^{(ss)}\right)}.$$

**$P53ser46$ equation at steady-state:**

$$\boldsymbol{k_d^{ser46}} = \frac{k_p^{ser46}\, P53^{(ss)}\, pATR^{(ss)}}{P53ser46^{(ss)}}.$$

For the sake of simplicity, $\mathrm{P53ser46^{(ss)}}$ was fixed equal to the 5% of $\mathrm{P53^{(ss)}}$.

**$mRNA_{MDM2}$ equation at steady-state:**

$$k_{td}^{MDM2} = \frac{k_t^{MDM2} + k_t^{MDM2P53}\dfrac{\left(P53^{(ss)} + P53ser46^{(ss)}\right)^{n_{P53}}}{(P53^{(ss)} + P53ser46^{(ss)})^{n_{P53}} + {K_{P53}}^{n_{P53}}}}{mRNA_{MDM2}^{(ss)}}.$$

$Mdm2$ **equation steady-state**

$$\boldsymbol{k_d^{MDM2}} = \frac{k_f^{MDM2} mRNA_{MDM2}^{(ss)}}{MDM2^{(ss)}}.$$

### 3. Step 2 of Model Parameter Estimation

#### 3.1 Parameter Estimation Strategy

In the *step 2* of model calibration, we estimated the remaining parameters by using a modified least square approach supplemented with semi-quantitative constraints. Numerical minimization was performed by Covariance Matrix Evolutionary Strategy, CMA-ES (Hansen and Ostermeier, 2001). At first, the initial parameter search interval was set to [0, 1] and initial value to 0.1 for all kinetics parameters, unless differently specified in Table S3. Next, the CMAES algorithm was run in iterative loops, the initial guesses being updated to the best solution found in the former loop and the search space being fixed as neighborhood of this optimal parameter set.

The model was calibrated using several experimental datasets available in both LN229 MGMT-sensitive and MGMT+ resistant cell lines. The biological studies integrated for parameter estimation are described hereafter:

-In Stratenwerth et al. *(10)*, O6-methyl-deoxyguanosine adducts were quantified for different TMZ doses after 3h of exposure (Fig. 3 (A)) and in a longitudinal manner during TMZ exposure (50 µM, unshown). For this last dataset, the actual TMZ dose was estimated to TMZout0_Ka_XTMZ as simulations suggested that the indicated dose was overestimated (Table S3).

-In He et al. *(21)* were published: dose-dependent $\gamma$H2AX foci assay after 72 h of TMZ exposure (Fig. 3(B)); TMZ dose-dependent western blot (WB) analysis of P53 and p-P53-Ser46 expression after 24 h and 72 h of exposure (Fig. 3(C,D)). As WB data are relative, we set the absolute value of the datapoint of P53 amount for the lowest dose of TMZ to be equal to the initial concentration of P53 (P53_0) as estimated in the model. For P53_ser46, we estimated an additional parameter, P53ser46_ref50_72, corresponding to the absolute value of P53_ser46 concentration after 72h of exposure to TMZ at 50 µM. This value was used to scale the other datapoints of P53_ser46 data to obtain absolute values, expressed in µM.

-In Jackson et al. *(18)*, several datasets were available including: cell survival after six days of exposure to TMZ at various concentrations (Fig. 3 (F)); time-dependent WB analysis of MGMT, CHK1 and pCHK1 expression levels after exposure to 12.5 µM or 100 µM of TMZ (Fig. 4 (C,I,J)). For CHK1 WB data, the average of the first three datapoints were set equal to the initial concentration of CHK1 (Chk1 _0), as estimated in the model. For pCHK1 WB data, we estimated an additional parameter, pCHK1_Jack_ref12_5_24h, corresponding to the absolute value of pCHK1 concentration after 24h of exposure to TMZ at 12.5 µM. This value was used to scale the other datapoints of pCHK1, to obtain absolute values, expressed in µM. Next, an additional dataset studying CHK1 and pCHK1 amount in control conditions or in the presence of TMZ and an inhibitor of ATR at various concentration was used. For CHK1, these WB values were normalized by setting equal the datapoint without drugs and Chk1 _0. For pCHK1, an additional parameter, pChk1_Jack_ATR_ref100_24h, was estimated corresponding to the absolute value of the datapoint at 24h, with TMZ (100 µM) and without any ATR inhibitor.

-In Aasland et al. 2020 *(17)*, we could retrieve: time-dependent WB analysis of pCHK1, CDC25, P53 and P53ser46 expression after exposure to 100 µM of TMZ (Fig. 4 (J, K, M, N)). CDC25 WB data were transformed into absolute values using initial protein concentrations estimated in the model, and an additional parameter, pChk1_Aas_ref100_48h, was calibrated to retrieve pCHK1 data in the correct unit. For P53, we used the shared conditions with the dataset from *(21)* (i.e. 24h of TMZ exposure at 100 µM, in MGMT- cells) to transform data into µM.

##### 3.2 Biological constraints applied to parameter estimation.

To ensure model accuracy and parameter identifiability, biological constraints were added to the procedure for parameter estimation. Briefly, in the minimization task performed by CMAES, the cost function was forced to explode (i.e. to take a very large value) whenever at least one of the constraints was not fulfilled, and the currently tested parameter set was discarded. Here are the constraints applied:

- **On $\boldsymbol{k_{addN}}$ and $\boldsymbol{k_{addO}}$:** from literature, it is known that N7-meG adducts accounts for 85% of all TMZ-induced base lesions, N3-meA/G for 10%, and O6-meG for 5% *(12, 26)*. To obtain the same percentage in our simulations, we accepted only the set of parameters for which the average of O6-meG concentration between 0 and 2h of exposure (i.e. the time frame with the O6-meG peak) was between 5/95 and 12/88 of the average of N37-meAG adduct over the same time window. For computational efficiency, we did not impose this constraint for all simulated cell liens and doses but only for TMZ=24$\mu$M in MGMT- cells (this was the scenario for which O6-meG amount was the greatest among all datasets, hence supposedly the one with the largest signal-to-noise ratio considering the low values of the species).
- **On $\boldsymbol{k_{addEx}}$:** we assumed that most of N37mAG adducts was generated by the TMZ-derived cation $C_{in}$ and only a small part by the TMZ-independent term $k_{addEx}$. Let $r$ be the ratio between the average of the term $k_{addN}DNAC_{in}$ in the first 6 hours (during which there is the peak of N37mAG) at TMZ=10$\mu$M (considered a low dose, but producing an amount of DNA adducts not neglectable) as numerator and $k_{addEx}$ as denominator, we impose $r \in [5{,}15]$.
- **On DSB concentrations**: we suppose that the maximum amount of DSB in MGMT+ cells could not overpass that in MGMT- cells during TMZ exposure. For the sake of computational efficiency, this constraint was written only for the dose TMZ=20$\mu$M for which we have the

greatest number of DSB data points. Thus, we only accepted the set of parameters for which the maximum of DSB in MGMT- was less than 1/3 of that in MGMT+, for TMZ=20$\mu$M. The fraction 1/3 has been chosen arbitrarily, according to the data.

- **On $\boldsymbol{K_{ATR}}$**: the value of the activation threshold, $K_{ATR}$, was approved only if it was between the smallest pATR simulated value, obtained with the dose TMZ=1 $\mu$M, and 1.5-fold the greatest one, obtained for TMZ=125$\mu$M. To simplify, we implemented this constraint only at time t=72h and related to the cell line MGMT-.
- **On $\boldsymbol{K_{p53}^{(cellpop)}}$:** a set of parameters was accepted only if P53 activation threshold, $K_{P53}$, was less than the maximum total amount of P53 obtained by simulations over 6 days, for the maximum dose of TMZ=1024 $\mu$M and for the cell line MGMT-.
- In the absence of TMZ, the cell cycle forcing function is supposed to remain unchanged, despite of the presence of low amount of DNA damage due to the term $k_{addEx}$. Hence, the function should behave as an oscillator with constant period, mean, amplitude and phase. According to the Cell cycle module equation, the only element that may be subject to change is the amplitude that may be decreased by the inhibition of CDC25. To prevent it when TMZ=0$\mu$M, we imposed a maximal variation of 5% between the maximum value of the S-phase marker (CCNA) on the first and fifth days of simulations.

7. I. C. Sorribes, S. K. Handelman, H. V. Jain, Mitigating temozolomide resistance in glioblastoma via DNA damage-repair inhibition. .

8. M. Xu-Welliver, A. E. Pegg, Degradation of the alkylated form of the DNA repair protein, O6-alkylguanine-DNA alkyltransferase. .

16. S. Caporali, S. Falcinelli, G. Starace, M. T. Russo, E. Bonmassar, J. Jiricny, S. D'Atri, DNA Damage Induced by Temozolomide Signals to both ATM and ATR: Role of the Mismatch Repair System. , 14.
