## Supplementary Figures and Tables for "Circumventing glioblastoma resistance to temozolomide through optimal drug combinations designed by quantitative systems pharmacology and machine learning"

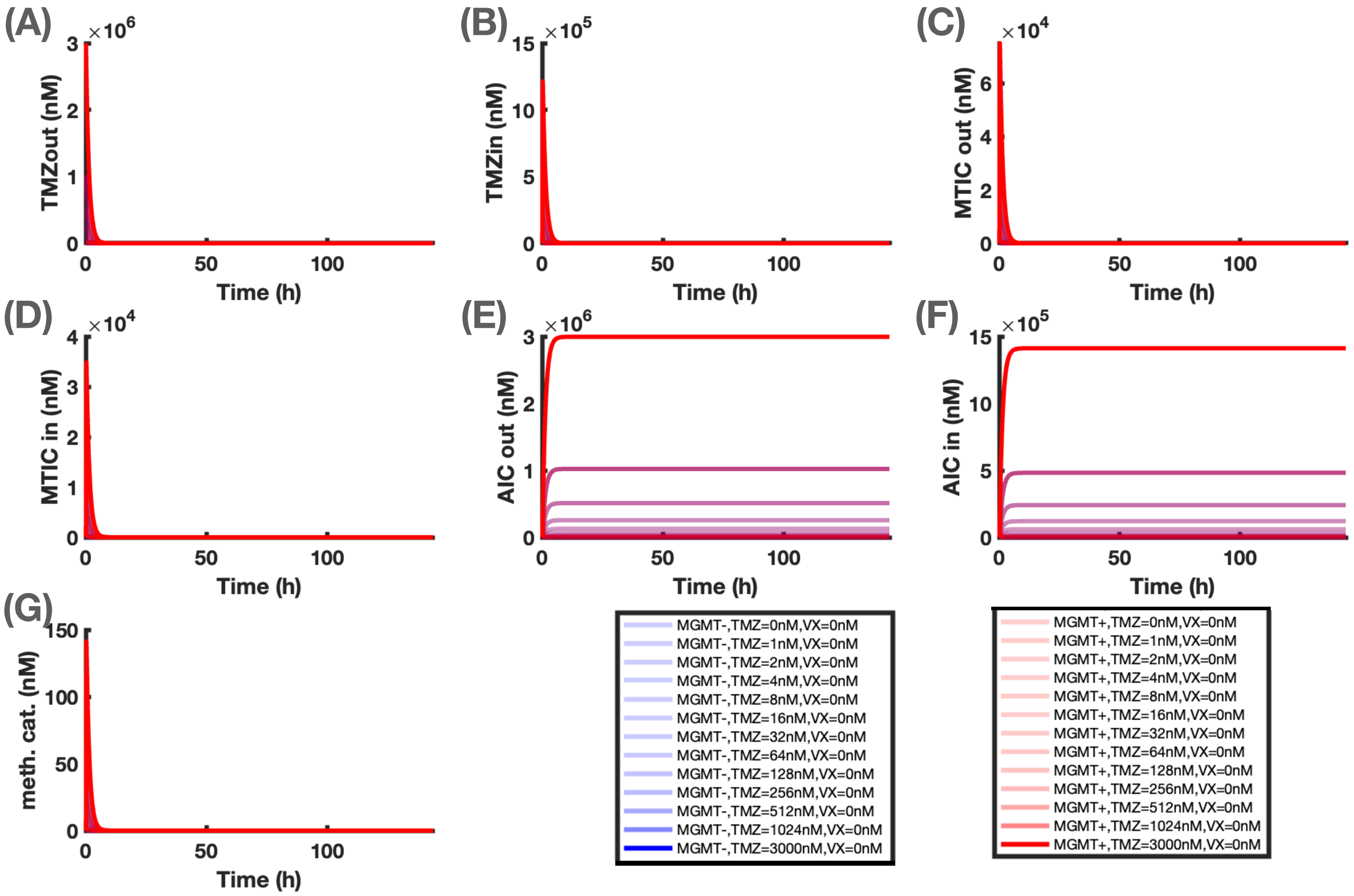


**Figure S1: Time-dependent TMZ cellular pharmacokinetics for indicated doses, in LN229 MGMT- and MGMT+ cells.**

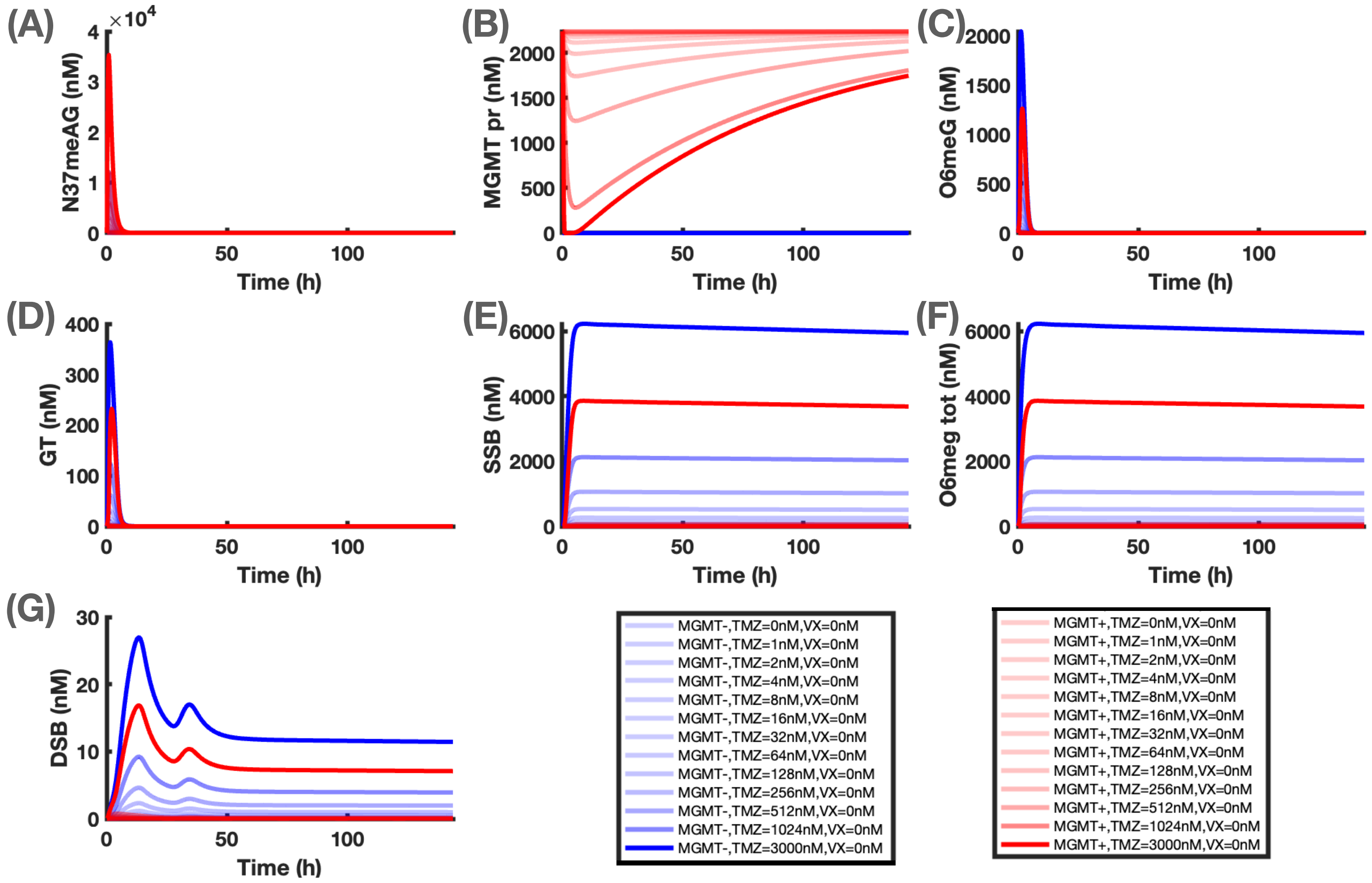


**Figure S2: Time-dependent TMZ-induced DNA damage, for indicated doses, in LN229 MGMT- and MGMT+ cells.**

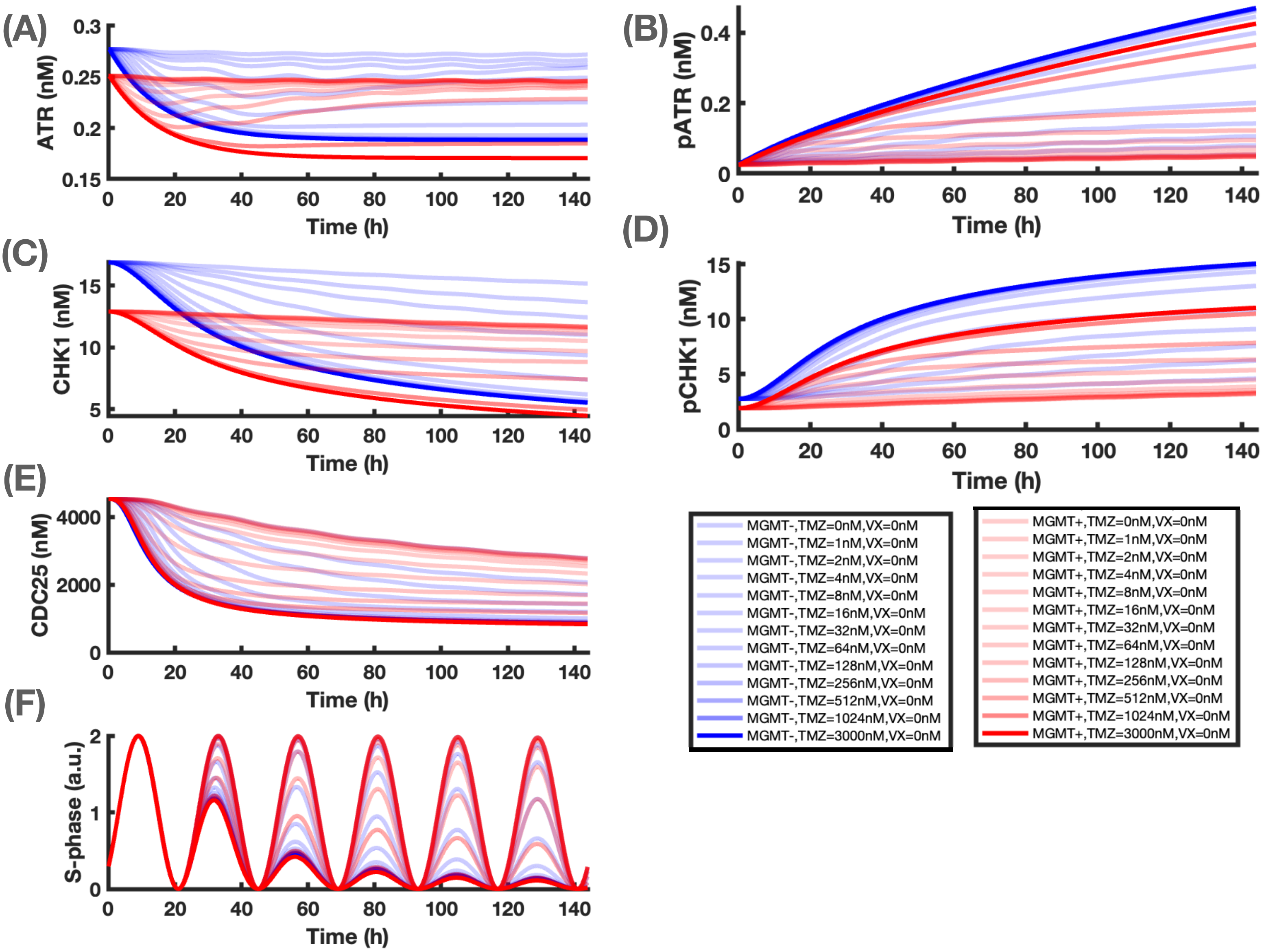


**Figure S3: Time-dependent TMZ pharmacodynamics and cell cycle arrest, for indicated doses, in LN229 MGMT- and MGMT+ cells.** VX refers to the ATR inhibitor VX970.

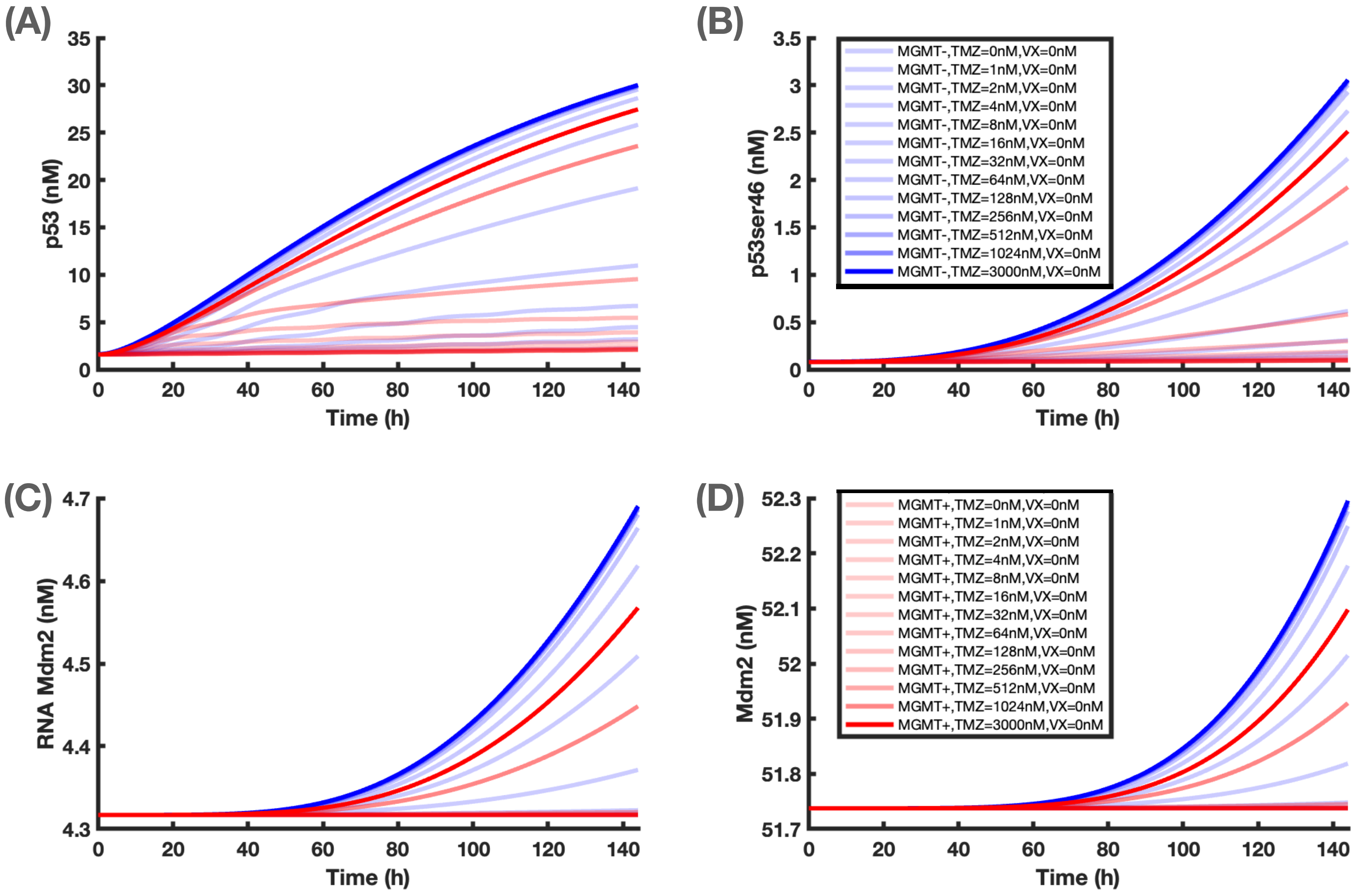


**Figure S4: P53-MDM2 network dynamics in the presence of TMZ at indicated doses, in LN229 MGMT- and MGMT+ cells.** VX refers to the ATR inhibitor VX970.

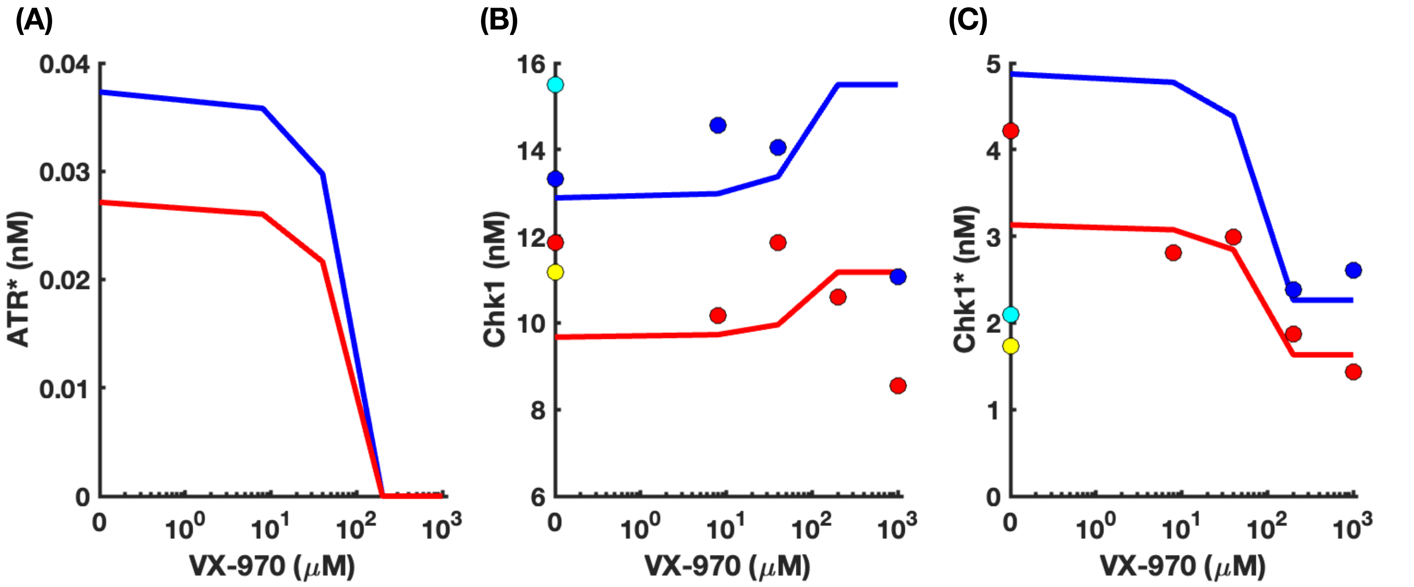


**Figure S5. ATR, Chk1 and pChk1 levels in MGMT- (blue) and MGMT+ (red) cells for various doses of the ATR inhibitor VX970 combined to TMZ, over 72h of exposure.** Dots are the data from Jackson et al. 2019 and solid curves are the best fit model.

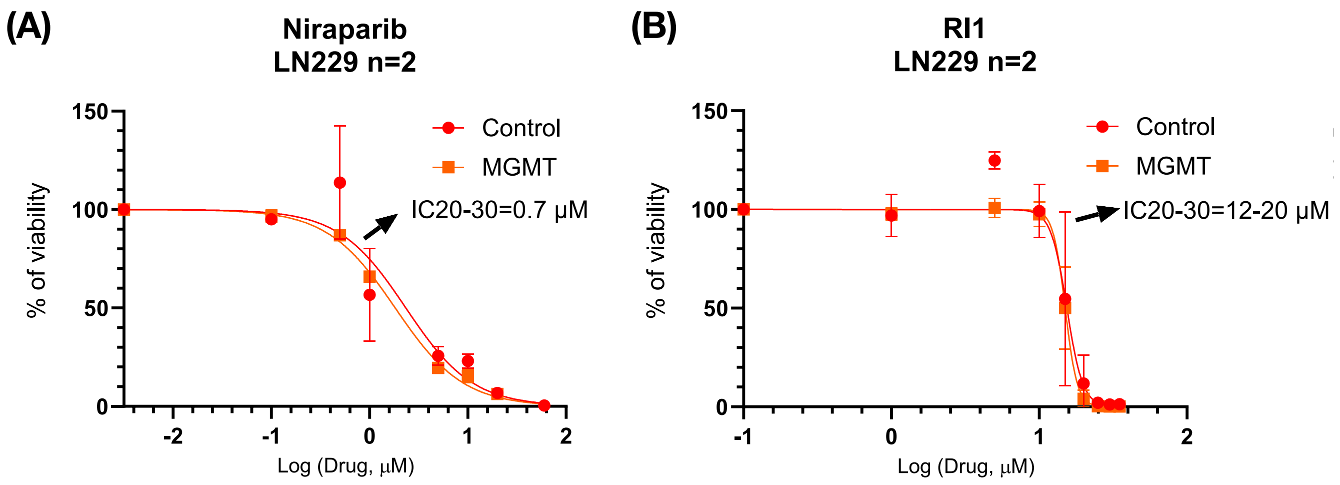


**Figure S6: Cell Viability after exposure to niraparib (BER inhibitor) (A) or RI-1 (HR inhibitor) (B) in MGMT- (red) or MGMT+ (orange) cells.**

### Supplementary Tables

| Variable | Biological meaning | Initial value in MGMT- cells (μM) | Initial value in MGMT+ cells (μM) | Reference |
|---|---|---|---|---|
| $T_{out}$ | TMZ extracellular concentration | TMZ_Dose | | Fixed as model input |
| $T_{in}$ | TMZ intracellular concentration | 0 | | SSE |
| $M_{out}$ | MTIC extracellular concentration | 0 | | SSE |
| $M_{in}$ | MTIC intracellular concentration | 0 | | SSE |
| $A_{out}$ | AIC extracellular concentration | 0 | | SSE |
| $A_{in}$ | AIC intracellular concentration | 0 | | SSE |
| $C_{in}$ | Methyldiazonium cation intracellular concentration | 0 | | SSE |
| $O6mG$ | O6-methylguanine DNA adducts | 0 | | SSE |
| $MGMT$ | MGMT protein | 0 | 2,238E+00 | Jackson et al. 2019 and estimated in step 2 |
| $GT$ | O6-meG:T mismatch | 0 | | SSE |
| $SSB$ | Single strand break | 0 | | SSE |
| $N37mAG$ | N7-methylguanine, N3-methylguanine and N3-methyladenine DNA adducts | 4,365E-02 | | SSE |
| $DSB$ | Double strand break | 9,856E-06 | | SSE and He et al. 2019 |
| $ATR$ | ATR protein | 2,772E-04 | 2,510E-04 | SSE |
| $pATR$ | phosphorylated ATR | 2,796E-05 | 2,532E-05 | SSE |
| $CHK1$ | CHK1 protein | 1,600E-02 | 1,289E-02 | SSE &Schwanhäusser 2011 |
| $pCHK1$ | Phosphorylated CHK1 | 2,770E-03 | 1,920E-03 | SSE |
| $CDC25$ | CDC25 protein | 4,527E+00 | | Schwanhäusser 2011 |
| $P53$ | P53 protein | 1,615E-03 | 1,500E-03 | SSE & Estimated |
| $P53ser46$ | P53 phosphorylated on serine 46 | 8,076E-05 | 7,871E-05 | 5% of P53 initial concentration |
| $mRNA_{MDM2}$ | MDM2 mRNA | 4,317E-03 | | Schwanhäusser 2011 |
| $MDM2$ | MDM2 protein | 5,173E-02 | | Schwanhäusser 2011 |

**Table S1. Model state variables and initial conditions.** SSE= steady-state equations, see Supplementary Information.

| Parameter Name | Biological meaning | Unit | Mean value | Standard deviation | Coefficient of Variation | Estimation | Data type | Reference |
|---|---|---|---|---|---|---|---|---|
| **TMZ PK** | | | | | | | | |
| **p_T** | TMZ cell uptake | $l\ h^{-1}$ | 3,70E-03 | 6,26E-04 | 1,69E+01 | | U87 | Ballesta 2014 |
| **p_T2** | TMZ cell efflux | $l\ h^{-1}$ | 7,79E-03 | 1,32E-03 | 1,70E+01 | | U87 | Ballesta 2014 |
| **k_T0** | TMZ activation into MTIC | $h^{-1}$ | 1,10E-07 | NA | NA | | Buffer solutions | Ballesta 2014 |
| **λ_T** | TMZ activation into MTIC | $h^{-1}$ | 2,09E+00 | 4,17E-04 | 2,00E-02 | | U87 | Ballesta 2014 |
| **k_M0** | MTIC transformation into AIC | $h^{-1}$ | 2,92E+02 | NA | NA | | Buffer solutions | Ballesta 2014 |
| **λ_M** | MTIC metabolism into AIC | $h^{-1}$ | 3,11E-01 | 8,08E-04 | 2,60E-01 | | U87 | Ballesta 2014 |
| **p_A** | AIC cell influx | $l\ h^{-1}$ | 3,56E-03 | 1,01E-03 | 2,85E+01 | | U87 | Ballesta 2014 |
| **p_A2** | AIC cell efflux | $l\ h^{-1}$ | 7,56E-03 | 2,15E-03 | 2,85E+01 | | U87 | Ballesta 2014 |
| **k_cat** | Methylating cation degradation | $h^{-1}$ | 6,00E+03 | NA | NA | | Buffer solutions | Ballesta 2014 |
| **DNA adduct formation** | | | | | | | | |
| **DNA** | Total number of DNA base pairs | μM | 5,50E+03 | NA | NA | | Human diploid cells | www.genome.gov |
| **k_addO** | O6-meG adduct formation rate | $h^{-1}$ | 1,12E-02 | 3,42E-04 | 3,04E+00 | | LN229 | |
| **k_addN** | N37-meAG DNA adduct formation rate | $h^{-1}$ | 1,54E-01 | 2,97E-02 | 1,93E+01 | | LN229 | |
| **kaddEx** | TMZ-independent N37-meAG DNA adduct formation rate | $\mu M\ h^{-1}$ | 9,43E-02 | NA | NA | D | LN229 | |
| **DNA repair** | | | | | | | | |
| **kf_MGMT_M ☆** | MGMT protein synthesis | $h^{-1}$ | 0,00E+00 | 0,00E+00 | NA | | LN229 MGMT- | Jackson 2019 |
| **kf_MGMT_P ☆** | MGMT protein synthesis | $h^{-1}$ | 1,76E+03 | NA | NA | D | LN229 MGMT+ | |
| **kd_MGMT** | MGMT protein degration | $h^{-1}$ | 1,10E-02 | NA | NA | | T98G | Smalley 2014 |
| **k_MGMT** | MGMT O6-meG adduct repair rate | $\mu M^{-1}\ h^{-1}$ | 4,76E+01 | 6,17E+00 | 1,30E+01 | | LN229 | |
| **k_BER** | N3meAG DNA adduct repair rate by BER (Base Excision repair) | $h^{-1}$ | 2,16E+00 | NA | NA | | Buffer solutions | Srivastava 1998 |
| **k_GT** | O6meGT formation rate | $h^{-1}$ | 1,00E+00 | NA | NA | Arbitrary | | |
| **k_MMR** | SSB formation rate from O6meG:T | $h^{-1}$ | 5,60E+00 | 1,04E+00 | 1,86E+01 | | HeLa | Geng 2011 |
| **k_DSBO** | DSB formation from SSBs | $h^{-1}$ | 3,18E-04 | 2,26E-05 | 7,12E+00 | | LN229 | |
| **k_DSBN** | DSB formation from N3/7meA/G | $h^{-1}$ | 3,75E-05 | 6,87E-06 | 1,83E+01 | | LN229 | |
| **k_HR** | DSB repair by HR | $h^{-1}$ | 1,66E-01 | NA | NA | | U87 | Short 2006 |
| **DNA damage response** | | | | | | | | |
| **kf_ATR_M ☆** | ATR protein synthesis rate | $\mu M\ h^{-1}$ | 1,22E-05 | NA | NA | | NIH3T3 mouse fibroblasts | Schwanhäusser 2011 |
| **kf_ATR_P ☆** | ATR protein synthesis rate | $\mu M\ h^{-1}$ | 1,10E-05 | 1,28E-07 | 1,16E+00 | | LN229 MGMT+ | |
| **k_ATR** | ATR protein activation rate | $h^{-1}$ | 2,11E-02 | 3,59E-03 | 1,70E+01 | | LN229 | |
| **n_add** | DSB Hill function exponential for ATR activation | | 2,00E+00 | NA | NA | Arbitrary | | |
| **K_add** | DSB Hill function threshold for ATR activation | μM | 7,00E-05 | 4,54E-06 | 6,49E+00 | | LN229 | |
| **kd_ATR** | ATR protein degradation rate | $h^{-1}$ | 4,35E-02 | NA | NA | | NIH3T3 mouse fibroblasts | Schwanhäusser 2011 |
| **kd_pATR** | Activated ATR degradation rate | $h^{-1}$ | 3,90E-03 | 7,06E-04 | 1,81E+01 | | LN229 | |
| **kf_Chk1_M ☆** | Chk1 protein synthesis rate | $\mu M\ h^{-1}$ | 1,75E-03 | 5,50E-04 | 3,15E+01 | | NIH3T3 mouse fibroblasts | Schwanhäusser 2011 |

| | | | | | | | | |
|---|---|---|---|---|---|---|---|---|
| **kf_Chk1_P ☆** | Chk1 protein synthesis rate | $\mu M\ h^{-1}$ | 1,32E-03 | 2,36E-05 | 1,79E+00 | | LN229 | |
| **k_Chk1** | Chk1 protein activation rate | $\mu M^{-1}\ h^{-1}$ | 4,88E+02 | 7,23E+01 | 1,48E+01 | | LN229 | |
| **kd_Chk1** | Chk1 protein degradation rate | $h^{-1}$ | 8,99E-02 | 8,46E-03 | 9,41E+00 | | NIH3T3 mouse fibroblasts | Schwanhäusser 2011 |
| **kd_pChk1** | Activated Chk1 degradation rate | $h^{-1}$ | 8,28E-02 | 1,13E-02 | 1,36E+01 | | LN229 | |
| **Cell Cycle** | | | | | | | | |
| **k_CCNA** | Cell cycle control rate | | 2,07E+00 | 2,99E-01 | 1,44E+01 | | LN229 | |
| **M_cc** | Cell cycle fixed magnitude | | 1,00E+00 | NA | NA | | | |
| **A_cc** | Cell cylce varible magnitude | | 1,00E+00 | NA | NA | | | |
| **T_cc** | Cell cycle period | h | 2,40E+01 | NA | NA | | | Gerard, 2009 |
| **phi_cc** | Cell cycle initial phase | h | 9,00E+00 | NA | NA | | | Gerard, 2009 |
| **n_cdc25** | Cell cycle Hill function exponantial for Cdc25 degradation | | 6,00E+00 | NA | NA | Arbitrary | | |
| **K_cdcd25** | Cell cycle Hill function threshold for Cdc25 degradation | $\mu M^{-1}$ | 1,37E+00 | 6,39E-02 | 4,67E+00 | | LN229 | |
| **n_cc** | Cell cycle control Hill function exponential | | 6,00E+00 | NA | NA | Arbitrary | | |
| **K_cc** | Cell cycle control Hill function threshold | $\mu M$ | 1,30E+00 | NA | NA | | | |
| **kf_cdc25_M ☆** | Cdc25 protein synthesis rate | $\mu M\ h^{-1}$ | 3,22E+00 | NA | NA | D | LN229 MGMT- | |
| **kf_cdc25_P ☆** | Cdc25 protein synthesis rate | $\mu M\ h^{-1}$ | 2,27E+00 | NA | NA | D | LN229 MGMT+ | |
| **k_cdc25** | Cdc25 protein inhibition rate | $\mu M^{-1}\ h^{-1}$ | 2,45E+02 | 6,08E+01 | 2,48E+01 | | LN229 | |
| **kd_cdc25** | Cdc25 protein degradation rate | $h^{-1}$ | 3,17E-02 | 6,56E-05 | 2,07E-01 | | NIH3T3 mouse fibroblasts | Schwanhäusser 2011 |
| **P53-MDM2 pathway** | | | | | | | | |
| **kf_p53** | p53 protein synthesis basal rate | $\mu M\ h^{-1}$ | 7,00E-04 | 1,00E-04 | 1,43E+01 | | LN229 | |
| **k_p53** | p53 protein synthesis rate from ATR | | 3,77E+01 | 3,64E+00 | 9,65E+00 | | LN229 | |
| **n_ATR** | ATR Hill function exponential for p53 activation | | 2,00E+00 | NA | NA | Arbitrary | LN229 | |
| **K_ATR** | ATR Hill function threshold for p53 activation | $\mu M$ | 4,00E-04 | 6,29E-05 | 1,57E+01 | | LN229 | |
| **kd_p53** | p53 protein degradation basal rate | $h^{-1}$ | 5,00E-01 | NA | NA | | SW480, SW620 | Hesse 2021 |
| **kd_p53Mdm2** | p53 protein degradation rate from Mdm2 | $h^{-1}$ | 1,44E+02 | NA | NA | | | Sturrock 2011 |
| **n_Mdm2** | Mdm2 Hill function exponential for p53 degradation | | 2,00E+00 | NA | NA | Arbitrary | | Sturrock 2011 |
| **K_Mdm2** | Mdm2 Hill function threshold for p53 degradation | $\mu M$ | 8,00E+00 | NA | NA | | | Sturrock 2011 |
| **kp_ser46** | p53 protein activation rate at ser46 | $\mu M^{-1}\ h^{-1}$ | 3,90E+00 | 5,04E-01 | 1,29E+01 | | LN229 | |
| **kd_ser46_M ☆** | p53ser46 degradation rate | $h^{-1}$ | 2,20E-03 | NA | NA | D | LN229 MGMT- | |
| **kd_ser46_P ☆** | p53ser46 degradation rate | $h^{-1}$ | 2,00E-03 | NA | NA | D | LN229 MGMT+ | |
| **kt_Mdm2** | Mdm2 RNA transcription basal rate | $\mu M\ h^{-1}$ | 1,01E-03 | 6,60E-05 | 6,51E+00 | | NIH3T3 mouse fibroblasts | Schwanhäusser 2011 |
| **kt_Md2m2p53** | Mdm2 RNA transcription rate from p53 | $\mu M^{-1}\ h^{-1}$ | 6,66E+00 | NA | NA | | | Sturrock 2011 |

| | | | | | | | | |
|---|---|---|---|---|---|---|---|---|
| **n_p53** | p53 Hill function exponential for Mdm2 traduction | | 4,00E+00 | NA | NA | | | Sturrock 2011 |
| **K_p53** | p53 Hill function threshold for Mdm2 traduction | µM | 5,38E-01 | 1,95E-01 | 3,63E+01 | | LN229 | |
| **ktd_Mdm2** | Mdm2 RNA degradation rate | $h^{-1}$ | 2,35E-01 | NA | NA | D | LN229 | |
| **kf_Mdm2** | Mdm2 protein synthesis rate | $h^{-1}$ | 4,97E-02 | 4,00E-03 | 8,05E+00 | | NIH3T3 mouse fibroblasts | Schwanhäusser 2011 |
| **kd_Mdm2** | Mdm2 protein degration rate | $h^{-1}$ | 4,10E-03 | NA | NA | D | LN229 | |
| **Apoptosis** | | | | | | | | |
| **k_apop** | apoptosis rate | $h^{-1}$ | 6,47E-02 | 4,03E-03 | 6,23E+00 | | LN229 | |
| **n_p53int** | p53 Hill function exponential for apoptosis | | 2,00E+00 | NA | NA | Arbitrary | LN229 | |
| **upAsy** | p53 Hill function threshold upper asymptote | µM | 2,56E-01 | 4,61E-02 | 1,80E+01 | | LN229 | |
| **tED50** | p53 Hill function threshold efficacy temporal median | $h^{-1}$ | 3,78E+02 | 6,86E+01 | 1,81E+01 | | LN229 | |
| **sness** | p53 Hill function threshold stiffness | $h^{-1}$ | 4,20E-03 | 6,77E-04 | 1,61E+01 | | LN229 | |
| **k_dis** | death cell disparition rate | $h^{-1}$ | 1,64E-02 | 2,11E-04 | 1,29E+00 | | LN229 | |

**Table S2. Best fit model parameter estimates.** A star in the first column indicates that different values were imposed for MGMT- and MGMT+ cell lines. In the "estimation" column, red indicates an estimation during step 1, blue, during step 2 and "D" indicates that the parameter was Derived from the steady state assumptions.

| Parameter | Range | Motivations |
|---|---|---|
| TMZout0_Ka_XTMZ | $[36,38]$ | According to our simulation, the set of four data points related to the expression of O6-meG and coming from *(25)* do not correspond to the TMZ dose of 50$\mu$M as indicated by the authors, but they seem related to a lower dose. At the beginning we imposed the range [25 $\mu$M,50 $\mu$M] but after several runs of the optimization algorithm, we adjusted the range at [36 $\mu$M.38 $\mu$M]. |
| MGMT_P0 | $[0.5,1.5]$ | The MGMT initial condition for the MGMT+ cell line is supposed to be greater than standard condition. For this reason, we imposed at the beginning to be greater than 100nM but lesser than 1500nM. After several runs of the optimization algorithm, we adjusted the range at [0.5 $\mu$M,1.5 $\mu$M]. |
| k_MGMT | $> 1$ | Range fixed after several runs of the optimization algorithm. |
| k_CCNA | $[1,20]$ | Range fixed after several runs of the optimization algorithm. |
| K_add | $[0.05e-3, 10e-3]$ | Range fixed after several runs of the optimization algorithm. |
| pCHK1_ref12_5_24h | $> 0.005\, *$Chk1_Jack_MGMT_M_norm_24h | We suppose than only a small but not negligible percentage of Chk1 is activated by the TMZ effect. Hence, we impose that value used as reference for the pChk1 data at TMZ=12.5$\mu M$ and time t=24h should be greater than the 0.5% of the corresponding Chk1 data (related to the cell line MGMT-). |
| pCHK1_Jack_ref1100_48h | $<$Chk1_Jack_inhATR_VXcon_MGMT_M_norm | Similarly, we impose that value used as reference for the pChk1 data at TMZ=100$\mu M$ and time t=24h should be lesser than the corresponding Chk1 data (related to the cell line MGMT-). |
| pCHK1_Ass_ref1100_48h | $<$Chk1_Jack_inhATR_VXcon_MGMT_M_norm | Similarly, we impose that value used as reference for the pChk1 data at TMZ=100$\mu M$ and time t=24h should be lesser than the corresponding Chk1 data (related to the cell line MGMT-). |
| K_CDC25 | $[0.6,0.8]\, *$cdc25c_Aas_norm_48h | To define a range and increase the identifiability, the Cdc25 activation threshold for the cell cycle arrest is imposed to be between the 60% and 80% of the data at TMZ=100$\mu M$ and time t=48h. In fact, for that dose and that time, the cell cycle is supposed to arrest. |
| K_ATR | $[0.01e-3, 0.15e-3]$ | The p53 data show a change of behavior between the doses of 20 $\mu M$ and 25 $\mu M$ of TMZ. To keep the same behavior for the model simulations, the ATR activation threshold for p53 should be between 1nM and 150nM. |
| P53ser46_ref50_72 | $[0.01,1]\, *$p53_He_hdTMZ_MGMT_M_72h_norm | We impose that the value used as reference for the p53ser46 data at time t=72h and TMZ=50$\mu M$ should be between the 1% and 100% of the corresponding p53 data. |
| K_P53 | $[0.001,0.011]$ | We impose that the p53 activation threshold for Mdm2 mRNA must be between 1nM and 11nM, which correspond to the initial condition used for p53. |
| upAsy | $[0,120]$ | Range fixed after several runs of the optimization algorithm. |
| sness | $> 0.010$ | This range has been fixed to increase the identifiability of both parameters upAsy and sness. |
| Initial cell population size | $[70,80]$ | According to the survival data at initial time |

**Table S3. Parameter search intervals.** See Supplementary Information for details.

**Cell line independent parameters**

| Parameter | Biological meaning | Unit | Mean | SD | %CV | Marginal Distribution | Parameters | Vine copula |
|---|---|---|---|---|---|---|---|---|
| pT | TMZ cell uptake | l $h^{-1}$ | 7,785E-03 | 1,323E-03 | 1,700E+01 | LogNorm | mean sd<br>-5,612 0,1678 | ✗ |
| pT2 | TMZ cell efflux | l $h^{-1}$ | 7,785E-03 | 1,323E-03 | 1,700E+01 | LogNorm | mean sd<br>-4,869 0,168 | ✗ |
| k_add0 | O6-meG adduct formation rate | $h^{-1}$ | 1,124E-02 | 3,420E-04 | 3,043E+00 | Normal model | mean sd<br>0,011 0,000342 | ✓ |
| k_addN | N37-meAG DNA adduct formation rate | $h^{-1}$ | 1,538E-01 | 2,973E-02 | 1,933E+01 | Skew Generalized Error model | mean sd nu xi<br>0,146 0,029 3,513 1,375 | ✓ |
| k_CyA | Cell cycle control rate | | 2,071E+00 | 2,988E-01 | 1,443E+01 | Skew Generalized Error model | mean sd nu xi<br>2,1895 0,297 2,682 1,110 | ✓ |
| K_cdc25 | Threshold for Cdc25 degradation | $\mu M^{-1}$ | 1,369E+00 | 6,388E-02 | 4,666E+00 | Skew Generalized Error model | mean sd nu xi<br>1,33668 0,06379 1,83066 0,78645 | ✓ |
| k_BER | DNA adduct repair rate by BER | $h^{-1}$ | 2,160E+00 | 4,320E-01 | 2,000E+01 | LogNorm | mean sd<br>0,750 0,198 | ✗ |
| k_MMR | SSB formation rate from O6meGT | $h^{-1}$ | 5,600E+00 | 1,040E+00 | 1,857E+01 | LogNorm | mean sd<br>1,705 0,184 | ✗ |
| k_DSBO | DSB formation from SSBs | $h^{-1}$ | 3,179E-04 | 2,264E-05 | 7,122E+00 | Normal model | mean sd<br>3,312e-04 2,264e-05 | ✓ |
| k_DSBN | DSB formation from N3/7meA/G | $h^{-1}$ | 3,747E-05 | 6,868E-06 | 1,833E+01 | Inverse Gaussian model | mean shape<br>4,037e-05 1,366e-03 | ✓ |
| k_HR | DSB repair by HR | $h^{-1}$ | 1,660E-01 | 3,320E-02 | 2,000E+01 | LogNorm | mean sd<br>-1,815 0,198 | ✗ |
| k_ATR | ATR protein activation rate | $h^{-1}$ | 2,110E-02 | 3,591E-03 | 1,702E+01 | InvGamma model | alpha beta<br>36,742 0,757 | ✓ |
| K_add | Threshold for ATR activation | μM | 7,000E-05 | 4,541E-06 | 6,487E+00 | Inverse Gaussian model | mean shape<br>7,451e-05 2,030e-02 | ✓ |
| k_Chk1 | Chk1 protein activation rate | $\mu M^{-1} h^{-1}$ | 4,875E+02 | 7,229E+01 | 1,483E+01 | Skew Generalized Error model | mean sd nu xi<br>456,602 71,949 2,242 1,485 | ✓ |
| kd_pChk1 | Activated Chk1 degradation rate | $h^{-1}$ | 8,280E-02 | 1,127E-02 | 1,361E+01 | Beta model | shape1 shape2<br>46,83 534,04 | ✓ |
| Cdc250 | Cdc25 protein initial condtion | μM | 4,527E+00 | 9,055E-01 | 2,000E+01 | LogNorm | mean sd<br>1,490 0,198 | ✗ |
| k_cdc25 | Cdc25 protein inhibition rate | $\mu M^{-1} h^{-1}$ | 2,448E+02 | 6,080E+01 | 2,483E+01 | Skew Generalized Error model | mean sd nu xi<br>255,573 62,104 4,473 36,252 | ✓ |
| kf_p53 | p53 protein synthesis basal rate | μM $h^{-1}$ | 7,000E-04 | 1,003E-04 | 1,433E+01 | Nakagami model | shape scale<br>1,103e+01 4,507e-07 | ✓ |
| k_p53 | p53 protein synthesis rate from ATR | $h^{-1}$ | 3,771E+01 | 3,640E+00 | 9,653E+00 | Skew Generalized Error model | mean sd nu xi<br>33,7589 3,6053 3,1673 0,8329 | ✓ |
| K_p53 | Threshold for Mdm2 traduction | μM | 5,375E-01 | 1,953E-01 | 3,634E+01 | Skew Generalized Error model | mean sd nu xi<br>0,6055 0,1962 2,4564 5,4565 | ✓ |
| kf_Mdm2 | Mdm2 protein synthesis rate | $h^{-1}$ | 4,970E-02 | 4,000E-03 | 8,048E+00 | LogNorm | mean sd<br>-3,00 4e-05 | ✗ |
| k_apop | apoptosis rate | $h^{-1}$ | 6,470E-02 | 4,032E-03 | 6,232E+00 | Generalized Error model | mean sd nu<br>0,064600 0,004042 2,506339 | ✓ |
| upAsy | p53 threshold upper asymptote | μM | 2,557E-01 | 4,609E-02 | 1,803E+01 | Skew Generalized Error model | mean sd nu xi<br>0,23243 0,04595 1,96161 1,55585 | ✓ |
| tED50 | p53 threshold efficacy temporal median | h | 3,782E+02 | 6,860E+01 | 1,814E+01 | Nakagami model | shape scale<br>7,652e+00 1,460e+05 | ✓ |
| sness | p53 Hill function threshold stiffness | $h^{-1}$ | 4,200E-03 | 6,771E-04 | 1,612E+01 | Nakagami model | shape scale<br>8,490e+00 1,587e-05 | ✓ |

**Cell-line specific parameters**

| | | | | | | | | |
|---|---|---|---|---|---|---|---|---|
| MGMT0_M | MGMT initial condition | μM | 0,000E+00 | 0,000E+00 | 0,000E+00 | | mean sd<br>0 0 | |
| MGMT0_P | | μM | 2,238E+00 | 1,235E-01 | 5,519E+00 | Skew Generalized Error model | mean sd nu xi<br>2,2971 0,1205 8,5122 0,1692 | ✓ |
| kf_ATR_M | ATR synthesis rate | μM $h^{-1}$ | 1,217E-05 | 2,434E-06 | 2,000E+01 | LogNormal | mean sd<br>-11,336 0,198 | ✗ |
| kf_ATR_P | | μM $h^{-1}$ | 1,102E-05 | 1,280E-07 | 1,162E+00 | Normal model | alpha beta<br>36,742 0,757 | ✓ |
| kf_Chk1_M | Chk1 protein synthesis rate ratio | μM $h^{-1}$ | 1,746E-03 | 5,590E-04 | 3,202E+01 | LogNormal | mean sd<br>-6,399 0,312 | ✗ |
| kf_Chk1_P = kf_Chk1_M *C_Chk1 | | μM $h^{-1}$ | 1,319E-03 | 2,364E-05 | 1,793E+00 | Skew Generalized Error model (C_Chk1) | mean sd nu xi<br>0,76163 0,01353 2,69889 1,10916 | ✓ |

**Table S4. Parameter distributions used for analysis of parameter importance.** In the last column, a green tick indicates that the distribution was estimated by vine copula. SD=standard deviation, CV= coefficient of variation.

**Cell-line independent parameters**

| Name | Biological meaning | Unit | Mean | Std | %CV | LogNorm Parameters |
|---|---|---|---|---|---|---|
| pT | TMZ cell uptake | $l\ h^{-1}$ | 3,704E-03 | 7,408E-04 | 20 | MU SIGMA<br>-5.617 0.198 |
| pT2 | TMZ cell efflux | $l\ h^{-1}$ | 7,785E-03 | 1,557E-03 | 20 | MU SIGMA<br>-4.875 0.198 |
| k_addO | O6-meG adduct formation rate | $h^{-1}$ | 1,124E-02 | 2,248E-03 | 20 | MU SIGMA<br>-4.507 0.198 |
| k_addN | N37-meAG DNA adduct formation rate | $h^{-1}$ | 1,538E-01 | 3,076E-02 | 20 | MU SIGMA<br>-1.891 0.198 |
| k_CyA | Cell cycle control rate | $h^{-1}$ | 2,0706 | 4,141E-01 | 20 | MU SIGMA<br>0.708 0.198 |
| K_cdc25 | Hill function threshold for Cdc25 degradation | $\mu M^{-1}$ | 1,369E+00 | 2,738E-01 | 20 | MU SIGMA<br>0.294 0.198 |
| k_MGMT | MGMT repair rate | $\mu M^{-1}\ h^{-1}$ | 4,760E+01 | 9,520E+00 | 20 | MU SIGMA<br>3.843 0.198 |
| k_BER | BER repair rate | $h^{-1}$ | 2,160E+00 | 4,320E-01 | 20 | MU SIGMA<br>0.750 0.198 |
| k_MMR | SSB formation rate from O6meGT | $h^{-1}$ | 5,600E+00 | 1,120E+00 | 20 | MU SIGMA<br>1.703 0.198 |
| k_DSBO | DSB formation from SSBs | $h^{-1}$ | 3,179E-04 | 6,358E-05 | 20 | MU SIGMA<br>-8.073 0.198 |
| k_DSBN | DSB formation from N3/7meAG | $h^{-1}$ | 3,747E-05 | 7,494E-06 | 20 | MU SIGMA<br>-10.211 0.198 |
| k_HR | HR repair rate | $h^{-1}$ | 1,660E-01 | 3,320E-02 | 20 | MU SIGMA<br>-1.815 0.198 |
| k_ATR | ATR protein activation rate | $h^{-1}$ | 2,110E-02 | 4,220E-03 | 20 | MU SIGMA<br>-3.878 0.198 |
| K_add | DSB-mediated ATR activation | μM | 7,000E-05 | 1,400E-05 | 20 | MU SIGMA<br>-9.569 0.198 |
| kd_pATR | Phosphorylated ATR degradation rate | $h^{-1}$ | 3,900E-03 | 7,800E-04 | 20 | MU SIGMA<br>-5.557 0.198 |
| k_Chk1 | CHK1 protein activation rate | $\mu M^{-1}\ h^{-1}$ | 4,875E+02 | 9,751E+01 | 20 | MU SIGMA<br>6.169 0.198 |
| kd_pChk1 | Phosphorylated Chk1 degradation rate | $h^{-1}$ | 8,280E-02 | 1,656E-02 | 20 | MU SIGMA<br>-2.510 0.198 |
| Cdc250 | CDC25 protein initial condition | μM | 4,527E+00 | 9,055E-01 | 20 | MU SIGMA<br>1.490 0.198 |
| k_cdc25 | CDC25 protein inhibition rate | $\mu M^{-1}\ h^{-1}$ | 2,448E+02 | 4,897E+01 | 20 | MU SIGMA<br>5.480 0.198 |
| kf_p53 | P53 protein basal synthesis rate | $\mu M\ h^{-1}$ | 7,000E-04 | 1,400E-04 | 20 | MU SIGMA<br>-7.281 0.198 |
| k_p53 | P53 protein ATR-induced synthesis rate | | 3,771E+01 | 7,542E+00 | 20 | MU SIGMA<br>3.610 0.198 |
| K_ATR | ATR-mediated p53 activation | μM | 4,000E-04 | 8,000E-05 | 20 | MU SIGMA<br>-7.792 0.198 |
| kp_ser46 | P53 protein activation rate | $\mu M^{-1}\ h^{-1}$ | 3,903E+00 | 7,806E-01 | 20 | MU SIGMA<br>1.342 0.198 |
| kf_Mdm2 | MDM2 protein synthesis rate | $h^{-1}$ | 4,970E-02 | 9,940E-03 | 20 | MU SIGMA<br>-3.021 0.198 |
| K_p53 | P53-mediated Mdm2 translation | μM | 5,375E-01 | 1,075E-01 | 20 | MU SIGMA<br>-0.640 0.198 |
| k_apop | apoptosis rate | $h^{-1}$ | 6,470E-02 | 1,294E-02 | 20 | MU SIGMA<br>-2.756 0.198 |
| upAsy | Apoptosis threshold upper asymptote | μM | 2,557E-01 | 5,114E-02 | 20 | MU SIGMA<br>-1.383 0.198 |
| tED50 | Apoptosis threshold median | h | 3,782E+02 | 7,564E+01 | 20 | MU SIGMA<br>5.915 0.198 |
| sness | Apoptosis threshold stiffness | $h^{-1}$ | 4,200E-03 | 8,400E-04 | 20 | MU SIGMA<br>-5.486 0.198 |

**Cell-line specific parameters**

| | | | | | | |
|---|---|---|---|---|---|---|
| MGMT0_M | MGMT protein initial condition | μM | 0,000E+00 | 0,000E+00 | 0 | MU SIGMA<br>0 0 |
| MGMT0_P | | μM | 2,238E+00 | 4,475E-01 | 20 | MU SIGMA<br>0.8010 0.0798 |
| kf_ATR_M | ATR protein synthesis rate | $\mu M\ h^{-1}$ | 1,217E-05 | 3,043E-06 | 25 | MU SIGMA<br>-11.336 0.198 |
| kf_ATR_P | | $\mu M\ h^{-1}$ | 1,102E-05 | 1,280E-07 | 25 | MU SIGMA<br>-11.435 0.198 |
| kf_Chk1_M | CHK1 protein synthesis rate ratio | $\mu M\ h^{-1}$ | 1,746E-03 | 4,365E-04 | 25 | MU SIGMA<br>-6.370 0.198 |
| kf_Chk1_P | | $\mu M\ h^{-1}$ | 1,318E-03 | 2,364E-05 | 25 | MU SIGMA<br>-0.300. 0.198 |

**Table S5. Distribution of parameters used for generating the virtual patient population.**
